# Accurate spliced alignment of long RNA sequencing reads

**DOI:** 10.1101/2020.09.02.279208

**Authors:** Kristoffer Sahlin, Veli Mäkinen

## Abstract

Long-read RNA sequencing techniques are establishing themselves as the primary sequencing technique to study the transcriptome landscape. Many such analyses are dependent on read alignments. However, the error rate and sequencing length of long-read technologies create new challenges for accurately aligning these reads. We present an alignment method uLTRA based on a novel two-pass collinear chaining algorithm. Furthermore, uLTRA can be used both as a stand-alone aligner and as a wrapper around minimap2 for improved alignments in gene regions. We show that uLTRA produces higher accuracy over state-of-the-art aligners with substantially higher accuracy for small exons on simulated and synthetic data. On biological data where true read location is unknown, we show several examples where uLTRA aligns to known and novel isoforms with exon structures that are not detected with other aligners. uLTRA is available at https://github.com/ksahlin/ultra.

## Introduction

The transcriptome has been identified as an important link between DNA and phenotype and is therefore analyzed in various biological and biomedical studies. For these analyses, RNA sequencing has established itself as the primary experimental method. Some of the most common transcriptome analyses using RNA sequencing data include predicting and detecting isoforms and quantifying their abundance in the sample. These analyses are fundamentally underpinned by the alignment of reads to genomes. As a transcriptomic read can contain multiple exons, alignment algorithms are required to handle split alignment of a read to multiple exonic regions of the genome, referred to as a *spliced alignment*.

Spliced alignment is a challenging computational problem, and a plethora of different alignment algorithms have been proposed for splice alignment of short-read RNA-seq, with some of the key algorithmic advances given in TopHat (Trapnell, Pachter, and Salzberg 2009), STAR (Dobin et al. 2013), HISAT (Kim, Langmead, and Salzberg 2015), GMAP (Wu et al. 2016), and HISAT2 (Kim et al. 2019). While short-read RNA sequencing has shown unprecedented insights into transcriptional complexities of various organisms, the read-length makes it difficult to detect isoforms with complicated splicing structure and limits quantification of isoform abundance (Zhang et al. 2017).

Long read transcriptome sequencing protocols such as Pacific Biosciences (PacBio) Iso-Seq sequencing (B. Wang et al. 2016) and Oxford Nanopore Technologies (ONT) cDNA and direct RNA sequencing (Workman et al. 2019) are now establishing themselves as the primary sequencing techniques to detect novel isoforms. Long-read sequencing technologies can sequence transcripts from end-to-end, providing the full isoforms structure and therefore offer accurate isoform detection and quantification. Such protocols have opened up the possibility to investigate the isoform landscape for genes with multiple gene copies (Sahlin et al. 2018) and complex splicing patterns (Tseng et al. 2019), as well as to accurately decipher allele (Tilgner et al. 2014) and cell specific (Gupta et al. 2018) isoforms. However, the long-read technologies also offer new algorithmic challenges because of the higher error rate and longer sequencing length which makes most short read alignment algorithms unsuitable for long read splice alignment (Križanović et al. 2018). Therefore, long transcriptomic reads have, similarly to short reads, prompted splice alignment algorithm development. Some short-read aligners have been modified for long-read splice alignment (Dobin et al. 2013; Wu et al. 2016), while other aligners designed for splice alignment of long reads (Li 2018; Marić et al., n.d.; Liu et al. 2019; Boratyn et al. 2019). A recent method also suggested improving long-read splice alignments using ensemble prediction of splice sites (Parker et al. 2021). First, splice sites present in the sample are predicted using an ensemble of reads aligned in the region. In a second step, reads are aligned again using the predictions as a guide. There are also methods for post-correction of long read splice alignments using ensemble-based predictions (Workman et al. 2019). However, methods that use alignments from multiple reads to form consensus splice site predictions may over-correct less abundant splice sites and other rare events. Due to this limitation, it is desired to have an accurate aligner that individually considers the best alignment for each read.

A particularly challenging task of long-read splice alignment is alignment to very small exons (<30nt). Firstly, because their length small exons can be highly repetitive in the genome and be shorter than the required seed match length of the aligner. Secondly, even if the size is larger than the minimum seed match length, a small exon is less likely to contain seed matches if there are errors present. The inability to align a read to small exons may cause downstream analysis tools to predict and quantify erroneous isoforms. In addition, we show in this study that splice aligners that use junction-specific alignment penalties can create spurious junctions by overfitting alignments to canonical splice sites such as GT-AG junctions.

To alleviate these limitations, we have designed and implemented a splice alignment algorithm uLTRA that aligns long-reads to a genome using an exon annotation. uLTRA uses a novel two-pass collinear chaining algorithm. In the first pass, uLTRA uses maximal exact matches (MEMs) between reads and the transcriptome as seeds, which is more robust and informative than a fixed-length seed approach employed by many seed-and-extend methods (Wu et al. 2016; Li 2018; Dobin et al. 2013; Marić et al., n.d.; Liu et al. 2019; Kent 2002). Candidate genes regions are then identified from the MEM chaining solution. In the second pass, we employ a second chaining algorithm. The second chaining algorithm in uLTRA allows approximate sequence matches and formulates a novel chaining problem that incorporates approximate matches, overlap, and gap costs into the formulation. The second pass also includes all the annotated exons of the candidate gene(s) including exons that did not have a MEM. This extra inclusion allows alignment to very small exons, and differ from other two-pass alignment methods such as deSALT (Liu et al. 2019) and Graphmap2 (Marić et al., n.d.). Since the core algorithm of uLTRA is designed for accurate alignments around annotated gene regions, it limits finding novel gene regions. However, uLTRA also includes a setting where it wraps around minimap2. In this setting, uLTRA uses minimap2’s primary alignments outside the regions indexed by uLTRA and chooses the preferred alignment of the two aligners in gene regions.

We demonstrate using controlled datasets that uLTRA, both as a stand-alone aligner and as a wrapper around minimap2, produces more accurate alignments than other aligners, particularly for small exons. We also use a dataset with ONT sequencing of synthetic SIRV transcripts (known isoforms) to demonstrate that uLTRA aligns more reads to transcripts that are known to be in the sample. Furthermore, we show on biological data from both PacBio and ONT that uLTRA aligns more reads to annotated isoforms and has alignments to more distinct isoform structures. Finally, we demonstrate that uLTRA produces alignments to known and novel isoform structures in the PacBio Alzheimer dataset that are not found by other aligners. These isoform structures come from genes that have been studied or linked to Alzheimer’s disease and motivate the utility of our method for a range of downstream analysis tasks such as isoform prediction and detection, splice-site analysis, isoform quantification and more. uLTRA is available at https://github.com/ksahlin/ultra.

## Results

### uLTRA Overview

uLTRA solves the algorithmic problem of chaining with overlaps to find alignments. The method consists of three steps. An overview of uLTRA is shown in Figure 1. We first construct subsequences of the genome referred to as *parts, flanks*, and *segments* (Fig 1A; details in Step 1 in Methods). This step is similar to the indexing step in other alignment algorithms, where the data structures do not need to be reconstructed for new sequencing datasets.

**Figure 1.**
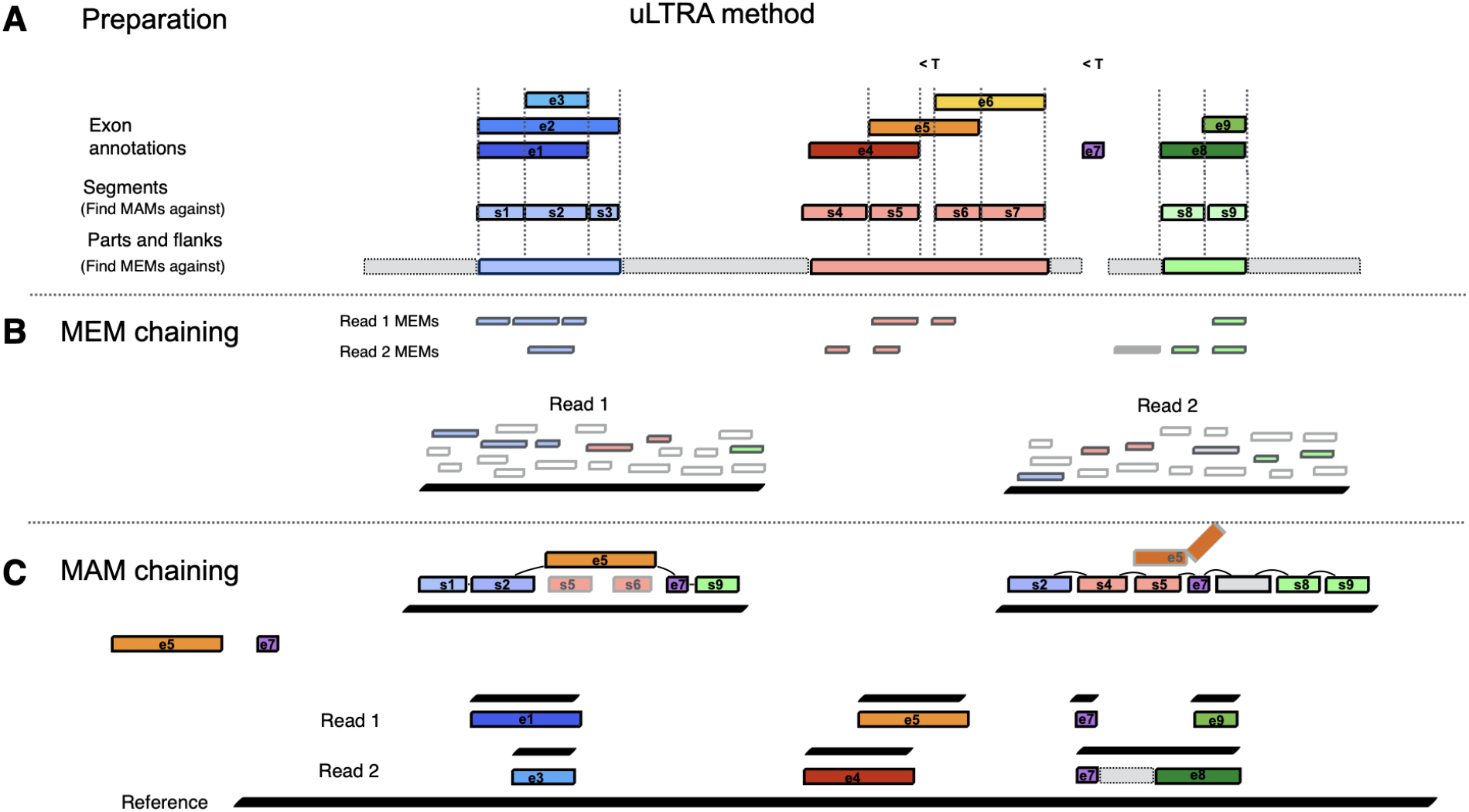
Overview of uLTRA alignment algorithm. (**A**) Segments, Parts and Flanks are stored and indexed for alignment. Small exons and segments below a threshold (indicated with < T in figure) are not indexed for MAM chaining but stored for the MAM chaining. (**B**) In the alignment step, MEMs in the reads to the parts and flanks are computed. Collinear chain(s) of MEMs covering as much of the read as possible are found for each read. (**C**) The solution consists of MEMs that overlap segments and/or flanks. These segments and flanks are linked to gene IDs. All the overlapping segments and flanks to the same gene IDs, including the small exons and segments excluded from the indexing, are retrieved and aligned to the read to form a set of MAMs. Collinear chains of MAMs are found by optimizing for coverage and alignment identity. The collinear chaining solution of MAMs is used to produce the final alignment of the read to the genome.

To align reads, uLTRA first finds maximal exact matches (MEMs) between the reads and the parts and flanks using slaMEM (Fernandes and Freitas 2014) (Fig. 1B). Each read will have a set of MEMs to the genome reference sequences (e.g., a set of chromosomes). Furthermore, we partition the instances within chromosomes if two consecutive MEMs on the chromosome are separated by more than a parameter threshold provided to uLTRA. For each instance, uLTRA finds a collinear chain of MEMs covering as much of the read as possible (allowing overlaps of MEMs in the read). We use Algorithm 1 in (Mäkinen and Sahlin 2020) to find such optimal chaining (see Step 2 in Methods). The optimal solutions to the instances produce candidate alignment sites.

In the third step, each solution to the MEM chaining is processed as follows. The MEMs in the chaining solution overlap distinct segments on the genome (segments defined in methods section; see Fig. 1B for illustration). Each segment belongs to a set of at least one gene. uLTRA aligns these segments together with all small exons (from the same genes) using edlib (Šošić and Šikić 2017). Each such alignment produces a maximal approximate match (MAMs; defined in methods section), and uLTRA uses all MAMs with alignment accuracy greater than a threshold *T* as input for the next chaining problem. There can be several MAMs of the same segment (or small exon) within a read. In the chaining of MAMs, we, roughly, optimize the total weight of MAMs covering the read while penalizing gaps and overlaps between MAMs. Here, weight is defined by the alignment accuracy and the length of the match (see Step 3 in Methods). The final set of MAMs produced from the optimal solution(s) constitutes a final set of segments on the genome (Fig. 1C). Finally, we align the final set of segments to the read using parasail (Daily 2016) (semi-global mode), which produces the final alignment(s) and cigar strings to the genome.

When uLTRA is used as a wrapper around minimap2, it runs minimap2 and parses minimap2’s alignments to find primary alignments outside the regions indexed by uLTRA. These alignments are not considered for alignment with uLTRA. uLTRA then proceeds to align all remaining reads. In a final step, uLTRA compares the reads that have been aligned with both aligners and selects the best alignment based on edit distance to the genome. The final output SAM-file consists of the best alignments to uLTRA-indexed regions and the alignments of minimap2 outside the regions indexed by uLTRA.

### Evaluation overview

We evaluated uLTRA against the two state-of-the-art transcriptomic long-read aligners minimap2 and deSALT. We also attempted to evaluate GraphMap2 but were unsuccessful in installing the aligner on one system and received segmentation fault on another (see supplementary note A). Several additional alignment methods can perform splice alignment of long transcriptomic reads such as BBMAP (Bushnell 2014), GMAP (Wu et al. 2016) STAR (Dobin et al. 2013), and HISAT2 (Kim et al. 2019). A recent benchmarking (Križanović et al. 2018) showed that GMAP performed the best among the tools compared on long noisy reads from complex genomes such as the human genome. Additional recent methods not included in (Križanović et al. 2018) include Graphmap2 (Marić et al. 2019), minimap2 (Li 2018), deSALT (Liu et al. 2019), and Magic-BLAST (Boratyn et al. 2019). However, in (Liu et al. 2019), the authors showed that deSALT, minimap2 outperformed GMAP across a large range of datasets, while in (Boratyn et al. 2019), which compared performance on both short and long reads, minimap2 performed the best for long noisy reads. Therefore, we compare uLTRA the more recent and best performing aligners minimap2 and deSALT. We run minimap2 and deSALT both with and without annotations as all three aligners support such modes. A tool that is run with annotations has ‘_GTF’ appended to its name. We also run uLTRA both as a stand-alone tool (labelled uLTRA) and as a wrapper around minimap2 (labelled uLTRA_mm2). Details for how the aligners were run are found in Supplementary Note A.

We used three *in silico*, one synthetic, and two biological datasets (Table 1) to evaluate the alignment algorithms. Of the biological datasets, two were from ONT and one from the PacBio Iso-Seq protocol. We used simulated datasets with known annotations to investigate the accuracy of spliced alignments as a whole, and of individual exons as a function of exon size. We used the synthetic SIRV data to investigate how aligners perform when aligning real sequencing reads to isoforms structures known to be in the sample. Finally, for the biological data where we do not have the ground truth annotations we measured the concordance in alignments between alignment methods. We also demonstrate that relying on alignment concordance as a proxy for alignment accuracy can be misleading due to similar alignment biases between aligners. We also report runtime and memory usage.

**Table 1.** Datasets included in evaluation. *Measured from minimap2’s alignments. Due to biological sequence variations, the error rate may be lower than the number presented here. ** Includes alternative haplotypes.

|  | Dataset | Nr reads | Median read length | Median error rate | Genome | Annotation |
| --- | --- | --- | --- | --- | --- | --- |
| Simulated | ENS | 234,207 | 890 | 0.0% | GRCh38.p12 | Gencode v34** |
|  | SIM_ANN | 1,000,000 | 864 | 8.6% | GRCh38.p12 | Gencode v34** |
|  | SIM_NIC | 1,000,000 | 1,272 | 8.6% | GRCh38.p1 | Gencode |
|  |  |  |  |  | 2 | v34** |
| Synthetic | SIRV | 1,514,274 | 538 | 6.9%* | SIRV genome | SIRV annotation C_170612a |
| Biological | ALZ | 4,277,293 | 2,699 | 1.2%* | GRCh38.p12 | Gencode v34** |
|  | DROS | 3,646,342 | 559 | 7.0%* | BDGP6.28 | Ensembl v100 |

### Alignment accuracy

We used three *in silico* datasets to test the alignment accuracy in a controlled setting (Table 1). First, we used 234,207 distinct cDNA sequences downloaded from ENSEMBL (denoted ENS) without introducing any simulated errors. We then simulated a dataset of 1,000,000 reads uniformly at random from the 234,207 ENSEMBL sequences with a mean error rate of 8.6% (denoted SIM_ANN for simulated annotated transcripts). Finally, to test the ability to align to transcripts containing novel combinations of exons, we simulated a dataset with the same error rate as SIM_ANN, that we call SIM_NIC for simulated Novel-In-Catalog transcripts. This dataset consists of reads from transcripts with novel exon combinations that we generated from gencode annotations (release 34, including haplotype scaffold annotations). See supplementary note B for details on the simulations. Since we have the true exon annotation of each read, we classify the read alignments as correct, inexact, exon difference, incorrect location, and unaligned. For details of these classifications, see Supplementary note B.

For SIM_ANN, which contains simulated reads from annotated transcripts, uLTRA has the highest fraction of correct alignments (93.6%) with a 2.8% percentage point increase compared to the second-best performing tool deSALT_GTF (Fig. 2A). uLTRA also substantially reduces errors classified as exon differences compared to the other aligners. Furthermore, we observed that uLTRA achieves considerably higher accuracy than other aligners for small exons (Fig. 2B). uLTRA_mm2 is further able to slightly increase accuracy over uLTRA (94.0%). For comparison, when minimap2 is run as a stand-alone tool, it has an accuracy of 87.9% (Fig. 2A). We observed similar trends for the ENS dataset (Fig. S1).

**Figure 2.**
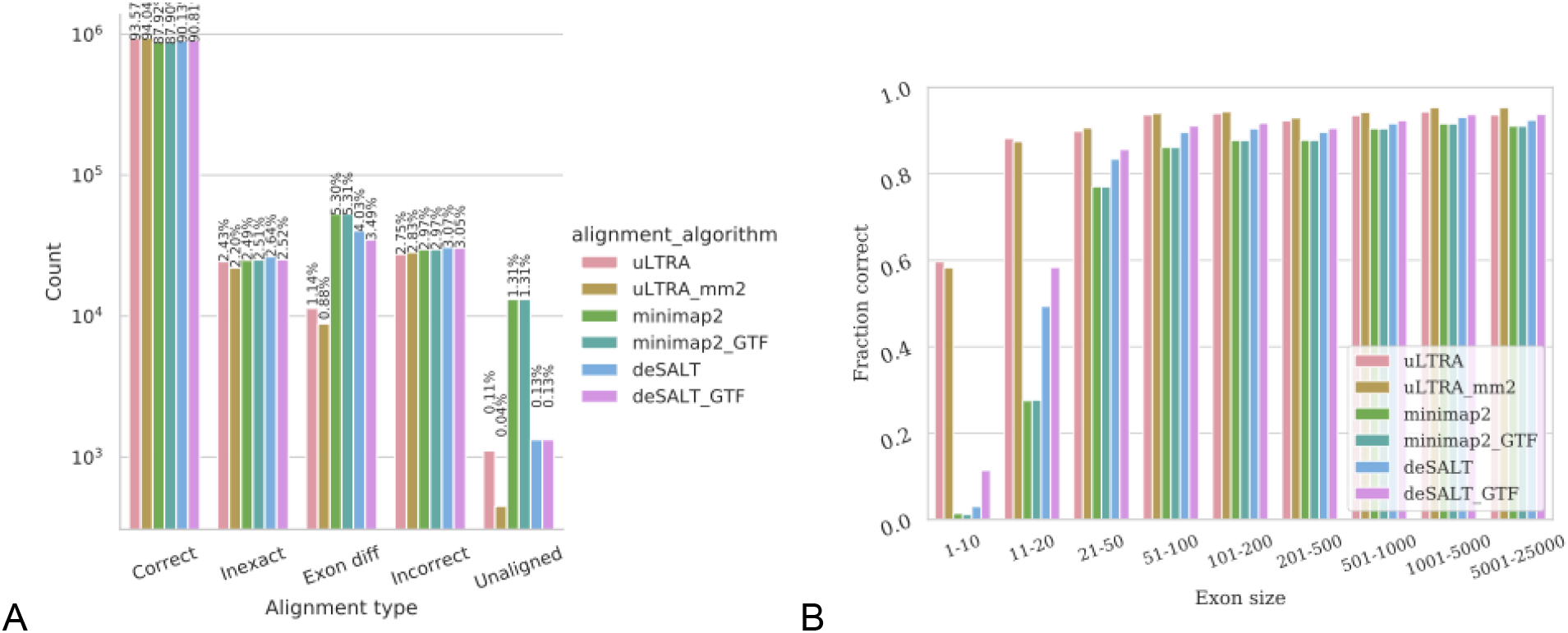
Alignment results on simulated data for the SIM_ANN dataset. (**A**) Percentage of reads in each respective category. (**B**) The fraction of correctly aligned exons (y-axis) as a function of exon size (x-axis).

As for the SIM_NIC, which contains only reads with novel combinations of exons, uLTRA’s accuracy is comparable to the ENS and SIM_ANN datasets (Fig. S2A). However, on this dataset uLTRA has a 9.6% percentage point more correctly aligned reads compared to the second-best performing aligner deSALT_GTF (Fig. S2A), and a 24.4% percentage point increase to minimap2. Our results show that the accuracy is substantially lower for the other aligners across exons sizes on this dataset (Fig. S2B), which disputes the explanation that the decrease in accuracy is caused solely by a larger fraction of smaller exons in the SIM_NIC dataset. The decreased accuracy may be explained by the fact that transcripts are simulated from all the reference sequences present in the Gencode release 34 (including haplotype scaffold annotations). This simulation will, therefore, produce more reads from transcripts with similar gene copies. As uLTRA’s accuracy remains similar to the other two datasets, it highlights the accuracy of aligning transcripts also to a reference genome with alternative haplotypes. Similarly to the other simulated datasets, uLTRA_mm2 has the highest accuracy (95.0%) which slightly improves over uLTRA as a stand-alone aligner.

### Splice site annotation performance on SIRV

We used the subset of 59 isoforms with distinct splice site positions from the ONT cDNA SIRV dataset (Sahlin and Medvedev 2021) to investigate alignment performance around splice sites (for details see Suppl. Note C). In this dataset, as the sequenced isoforms are known, we have a complete isoform annotation. We computed all alignments that had perfect matching splice sites to the annotations and denoted these reads as Full Splice Matches (FSM) following the notation in (Tardaguila et al. 2018). With the SIRV dataset, we have the properties of real ONT sequencing errors and genes, each expressing several known isoforms. The downside with SIRV data is that it does not represent the sequence complexity of a genome. For minimap2 and deSALT, we used non-default alignment parameters not to penalize non-canonical splice sites as much as in biological data. After this modification, we observed substantially improved alignment performance over default parameters (for details, see Suppl. Note A).

While sequencing bias may distort the read coverage per isoform and produce a dataset with different coverage distribution to what is present in the sample, the E0 mix contains transcripts at roughly equal abundances. In large, we observe similar distribution in the number of FSM alignments per isoform (Fig. S3) for all the aligners, but minimap2, minimap2_GTF, and deSALT produce less FSM alignments compared to deSALT_GTF, uLTRA, and uLTRA_mm2 across most isoforms. A notable difference is that minimap2 only aligns FSM reads to 54 unique isoforms, even after employing specific alignment parameters for SIRV data (Suppl. Note A). In comparison, deSALT and uLTRA in both settings were able to align FSM reads to all 59 unique isoforms.

There are some exceptions that we will now discuss. Firstly, both deSALT and minimap2 in annotation-free and annotation-provided settings align substantially fewer reads to SIRV503 than uLTRA and uLTRA_mm2. SIRV503 contains an 8nt long exon that deSALT and minimap2 do not align to in the large majority of reads containing the exon. Secondly, minimap_GTF does not align any reads to SIRV511, SIRV708, SIRV304, SIRV403, and SIRV408, while the annotation-free version of minimap2 does. Instead, we can see that minimap_GTF aligns a substantially larger fraction of reads to, e.g., SIRV506. Without setting specific alignment parameters for this dataset (Suppl. Note A), we observed that deSALT_GTF did not align any reads to SIRV511 and SIRV708. Such cases show that using specific alignment parameters for non-canonical splice sites may introduce alignment bias from overfitting to specific isoforms. Thirdly, comparing the two reference-free alignment methods minimap2 and deSALT, we observe that deSALT produces substantially fewer FSM alignments to minimap2 across most isoforms.

Overall, we observed more evenly distributed FSM alignment across the SIRV isoforms with both uLTRA and uLTRA_mm2. While there is no ground truth for this dataset, an equal abundance of isoforms is expected in this dataset from the design of the SIRV E0 mix. Furthermore, the number of FSM isoforms between uLTRA and uLTRA_mm2 alignments stays consistent.

### Evaluating alignments on biological data

We used an Alzheimer brain Iso-Seq dataset (denoted ALZ) and an ONT cDNA sequencing dataset from Drosophila (Sahlin and Medvedev 2021) (denoted DROS). Both datasets have been processed with respective bioinformatics pipelines to select only the reads containing full-length transcripts (for details see Suppl. Note D).

We neither have the correct read annotations, nor are we guaranteed to have a complete gene annotation for the biological datasets, which presents a challenge when evaluating accuracy. We took the following approaches. We first compared the aligned read categories between alignment algorithms following the definitions in (Tardaguila et al. 2018) (presented in the next section). Secondly, we looked at the alignment concordance between methods. Here we investigated concordance with respect to both alignment location on the genome and concordance in the categories of reads. Thirdly, we provide several examples of uniquely detected isoforms by uLTRA (and uLTRA_mm2), which demonstrate that alignment concordance analysis without ground truth has caveats.

### Alignment categories on biological data

We evaluated alignment categories to the previously annotated database using the categories defined in (Tardaguila et al. 2018). As in (Tardaguila et al. 2018), we classify an alignment of a read to the genome as a Full Splice Match (FSM), Incomplete Splice Match (ISM), Novel In Catalog (NIC), Novel Not in Catalog (NNC), or NO_SPLICE. An FSM alignment means that the combination of splice junctions in the read alignment has been observed and annotated as an isoform. An ISM alignment means that the combination of splice junctions is in the annotation, but it is missing junctions compared to the annotated models in either the 3’ or 5’ end. A NIC alignment consists of junctions that all appear in the annotation, but not together in a single isoform. An NNC alignment means that the read aligns with at least one junction that is not in the annotation, while NO_SPLICE are all alignments without splice sites. See (Tardaguila et al. 2018) for details regarding these definitions.

In (Tardaguila et al. 2018), the authors noted that a higher fraction of isoforms represented by FSM and NIC reads could be validated using orthogonal techniques compared to the NNC reads, where the large majority could be validated and may stem from sequencing artifacts or misalignments. As we neither know the true isoforms present in the samples nor have a complete annotation of all isoforms on the genome, comparing the alignment categories between alignment methods does not evaluate alignment performance. Nevertheless, the categories can be compared between aligners for general insight on alignment concordance. Furthermore, these alignment categories are important for various downstream isoform detection and classification methods such as SQANTI (Tardaguila et al. 2018), TAMA (Kuo et al. 2020), or TALON (Wyman et al., 2019). Therefore, we present the results here.

Overall, the aligners and their different modes produce a similar distribution of the different alignment categories on both the DROS and ALZ datasets (Fig. 3). We observe that uLTRA, uLTRA_mm2 and deSALT_GTF align more FSM reads than deSALT, minimap2 and minimap2_GTF. In the ALZ dataset, uLTRA has many unaligned reads due to a large fraction of reads (17.6%) aligning outside uLTRA indexed regions. We observe that the other aligners, including uLTRA_mm2, have no unaligned reads. Instead, they attribute a larger fraction of reads in the category NO_SPLICE (Fig. 3B). It is known that a substantial fraction of reads in long-read transcriptome sequencing data is coming from so-called intra-priming reads (Tardaguila et al. 2018). These reads are characterized by aligning without splice junctions to an unannotated genome location that contains a poly-A stretch downstream from their 3’ end. While not fully characterized, these reads are likely to be artifacts in the sequencing protocol and often filtered out in downstream analysis (Tardaguila et al. 2018).

**Figure 3.**
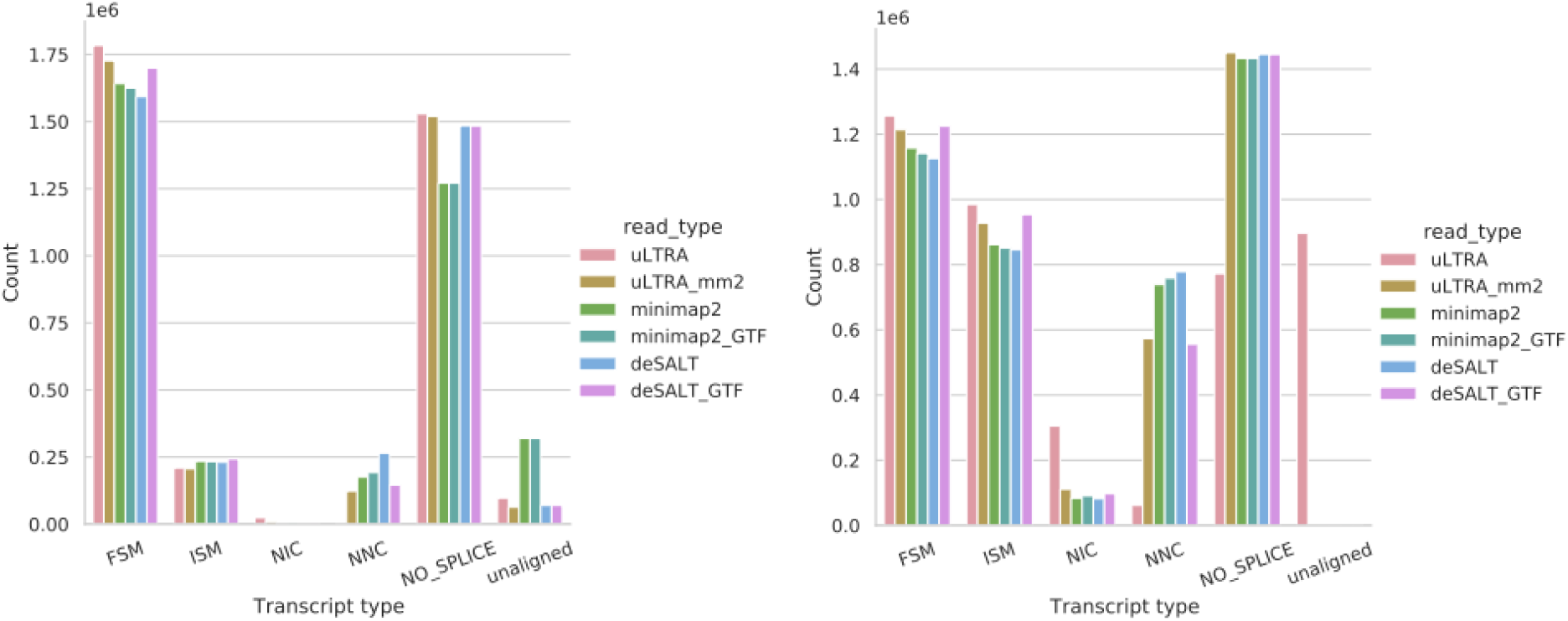
Number of reads annotated in different splicing categories for DROS (A) and ALZ (B).

As for the number of unique isoforms aligned to in the ALZ dataset, uLTRA has FSM alignments to more unique isoforms (39,384) compared to uLTRA_mm2 (36,776), deSALT_GTF (36,244), minimap2 (34,831), minimap2_GTF (34,495), and deSALT (34,421). We observed this trend also in the DROS dataset, where uLTRA had FSM alignments to 13,950 unique isoforms compared to uLTRA_mm2 (13,464), deSALT_GTF (13,367), minimap2 (13,092), minimap2_GTF (13,039), and deSALT (12,847). uLTRA also aligns a higher fraction of NIC reads in both the DROS and ALZ datasets (Fig. 3). While there is no ground truth for the datasets, likely, the substantial increase in unique isoforms by uLTRA as a stand-alone tool comes from the inability to align reads to genomic and pseudo-gene regions. Therefore, it is beneficial not to be limited to alignments around gene regions when aligning datasets where genomic contamination or novel gene regions exist.

We further investigated concordance in alignments between uLTRA_mm2, deSALT_GTF, and minimap2. They represent the best setting for each aligner, respectively, based on our accuracy evaluation on simulated data (Fig. 2, Fig. S1-2) and alignment consistency analysis on the synthetic SIRV data (Fig. S3).

### Alignment concordance on biological data

We first looked at a relaxed measure of alignment concordance. We define a read to have *concordant alignments* between two methods if the two alignments have a non-zero overlap on the genome (based on the start and stop coordinates). Note that this definition only captures discordance if the read aligns to different genes, not smaller differences around exons. A caveat with this definition occurs when measuring alignment concordance between more than two aligners. An alignment spanning positions A to C in one alignment may overlap with two disjoint alignments A to B and B+1 to C. In this case, we treat all the alignments as discordant. Finally, there are genes with multiple identical copies on the genome. In these cases, the alignment methods may choose different alignment locations simply by randomly picking a location. With these limitations in mind, we observed 90.3% and 98.6%% of all aligned reads had concordant genomic positions in DROS and ALZ, respectively (Fig. S4). This indicates that the mapping region is largely consistent between aligners and that most of the variability occurs in alignments around exons. The lower alignment concordance for the DROS dataset may result from a higher median error rate combined with a shorter average read length. In the DROS dataset, the second-largest category was the alignment concordance between uLTRA_mm2 and deSALT_GTF (5.7%; Fig. S4A).

We also looked at alignment concordance broken down individually within the classes FSM, ISM, NIC, NNC, and NO_SPLICE. For the FSM, ISM, NIC classes where splice sites are known, we classify an alignment as concordant between aligners if all the splice sites are identical. For NNC and NO_SPLICE, we use genomic overlap as described above. We observed a large concordance of alignments in the categories FSM, ISM, and NO_SPLICE and a slightly lower concordance for the categories NIC and NNC in both DROS (Fig. S5) and ALZ (Fig. S6) datasets. However, the NIC and NNC categories contain fewer reads (Fig. 3).

Finally, we also looked in more depth at the concordance of unique isoforms that had FSM predictions (Fig. S7). We observed a large concordance in the predicted FSMs. In total, 93.6% and 90.1% of the total unique isoforms with FSM alignments were aligned to by all the three methods on both datasets, for DROS and ALZ, respectively. The second largest category was isoforms aligned to by both uLTRA_mm2 and deSALT_GTF (2.2% and 4.3% for DROS and ALZ respectively).

### Caveats of assessing alignments based on concordance

We further investigated some of the isoform structures uniquely aligned to by uLTRA_mm2 and observed that in several cases, uLTRA’s alignments were correct. This analysis highlights that alignment concordance between aligners may not indicate correct alignment as concordance can come from the same algorithmic decisions between aligners such as customized alignment penalties for canonical regions or inability to align to very short exons.

#### FSM isoforms uniquely found by uLTRA

On the DROS dataset, uLTRA_mm2 aligned 338 FSM reads to 104 distinct isoforms (0.8% of total distinct isoforms) that minimap2 and deSALT_GTF did not align to. Of these isoforms, 7 had more than ten reads aligned, while most other isoforms had a coverage of 1-10 reads (Fig. S8A). For the ALZ dataset, uLTRA aligned a total of 9,130 FSM reads to 571 distinct isoforms (1.6% of total distinct isoforms) that minimap2 and deSALT_GTF did not align to. A total of 109 of these isoforms had more than 10 reads aligned, while most other isoforms had a coverage of 1-10 reads (Fig. S8).

We manually inspected a subset of the more abundant uniquely predicted isoforms by uLTRA_mm2 for the ALZ dataset using IGV (Robinson et al. 2011). We observed that some of these isoforms contained small exons (<10nt) that after manual inspection appeared correctly aligned to (Fig. S9). However, deSALT_GTF and minimap2 agreed on a different splicing structure, with the small exons put as an insertion or substitutions in the 5’ or 3’ ends of upstream or downstream exons. These alignments would show up as concordant between deSALT_GTF and minimap2_GTF in our previous analysis, although they are unlikely to be correct. The isoforms in Figure S9 come from the genes AP2, APBB, HNRNPM, and DCTN2, which come from gene families that have appeared in studies related Alzhemers’s disease (Tian et al. 2013) (Tanahashi and Tabira 1999) (Geuens, Bouhy, and Timmerman 2016) or other neurodegenerative disorders (Boland et al. 2018). All of these genes are supported by more than 100 reads and have perfect alignment across the junctions in uLTRA_mm2’s alignments.

In addition, we highlight another case of a potential subtle misalignment (Fig. S10) that makes the best fit FSM isoform go undetected. This potential misalignment is caused by using GT-AG specific alignment penalties and causes 500 reads to support a GT-AG splice junction in deSALT_GTF and minimap2. In this example, uLTRA_mm2’s alignments support a GC-AG junction. While we have no ground truth, and an insertion of one nucleotide near the splice site is plausible, uLTRA_mm2’s alignments best fit the data (omitting prior belief of GT-AG junction) and also supports a previously annotated isoform. The PRNP gene has also been studied in Alzheimer’s disease (Bagyinszky et al. 2019).

Finally, we illustrate an example (Fig. S11) of an instance of 161 reads where all three aligners have discordant alignments, caused by a segment of 9nt from a transcript from the SPOCK gene, which has also appeared in studies on neurodegenerative disorders (Charbonnier et al. 1997). Here, uLTRA_mm2 and deSALT_GTF align the 9nt portion of the read corresponding to two different exons while minimap2 does not align this region. Both uLTRA_mm2 and deSALT_GTF alignments are FSM but to different isoforms, and both upstream and downstream junctions are GT-AG. With this information, it is ambiguous as to which alignment is the correct one.

#### NIC isoforms uniquely found by uLTRA

We also observed that uLTRA_mm2 aligns more reads as NIC alignments compared to deSALT_GTF and minimap2 (Fig. 3, Fig. S5C, and S6C). Many of these NIC reads could be spurious due to inaccuracies in uLTRA_mm2 alignments when both upstream and downstream flanks of the junction contain the same nucleotide. We looked at the most abundant NIC uniquely aligned to by uLTRA_mm2, a transcript from the MBP gene (Fig. S12; predicted by 943 reads). uLTRA aligned reads to this NIC because of the homopolymer length difference of C’s in the reads, together with that both the upstream and downstream junction contained C’s. However, deSALT_GTF and minimap2 always aligned to the CT-AC junction by creating insertions of C at downstream junctions if needed (matching an FSM), while uLTRA_mm2 chooses a CT-TA junction for the reads where the homopolymer length was four cytosine nucleotides (creating a NIC). It is ambiguous as to what is the correct isoform in this example.

As for the uniquely predicted FSMs, the NICs also contained predictions with small exons (in total 50 unique NIC isoforms have exons smaller than 20nt). We manually inspected some of the more abundant predictions of which, similarly to the FSMs with small exons, the data supports their correctness (Fig. S13 A-C). The three isoforms presented in Fig. S13 are highly supported isoforms from the MICU1, SEPTIN7, and APBB1 genes and are, furthermore, novel with respect to the Gencode v34 annotation and have appeared in studies on Alzheimer’s disease (X. Wang et al. 2018) (Calvo-Rodriguez et al. 2020) (Tanahashi and Tabira 1999).

### Runtime and memory usage

We used a 128Gb memory node with 20 cores. We tested the tools using both 4 and 19 cores (leaving one core for the main process) to study parallelization performance. We measured user time (total time from start to finish) and peak memory usage (highest memory usage across the program lifetime).

Using 4 cores, we observe that deSALT is the fastest tool except on the largest dataset (ALZ) where minimap2 is the fastest (Table 2). The relative runtime difference between uLTRA and deSALT decreases with the organism’s size and the number of reads on our datasets. For example, on the two smaller datasets SIRV and DROS, deSALT is about 6 times and 4 times faster, respectively. While on the largest datasets, SIM_NIC and ALZ, deSALT is about 2 times and 1.6 times faster, respectively (Table 2). uLTRA has a similar or faster runtime than minimap2 on ENS, SIM_ANN, and SIM_NIC, respectively, but is slower on the other datasets. Using uLTRA as a wrapper around minimap2 increases the runtime slightly, but it is substantially less costly than running both aligners separately. For the ALZ dataset, running uLTRA_mm2 is only about 8% slower than only running uLTRA. Overall, while uLTRA and uLTRA_mm2 have a slightly larger runtime than minimap2 and deSALT, the practical difference, particularly for the larger datasets, is not major.

**Table 2.** Runtime of alignment using 4 cores.

| Dataset | uLTRA | uLTRA_mm2 | minimap2 | minimap2_GTF | deSALT | deSALT_GTF |
| --- | --- | --- | --- | --- | --- | --- |
| ENS | 52m | 1h 11m | 43m | 45m | 18m | 17 |
| SIM_ANN | 2h 47m | 4h 00m | 2h 42m | 2h 48m | 1h 20m | 1h 23 |
| SIM_NIC | 3h 21m | 6h 40m | 4h 35m | 4h 42m | 1h 46m | 1h 55m |
| SIRV | 35m | 50m | 13m | 13m | 6m | 7m |
| ALZ | 16h 9m | 17h 32m | 9h 4m | 9h 47m | 10h 16m | 10h 17m |
| DROS | 1h 17m | 1h 37m | 18m | 25m | 23m | 23m |

When we supply 19 cores for alignment, we see similar trends as to using 4 cores. The alignment time difference between uLTRA and the other aligners is the largest for smaller datasets such as SIRV and DROS, and evens out for larger datasets SIM_NIC and ALZ (Table S1). For example, uLTRA is about 80% slower on the ALZ dataset than both minimap2 and deSALT.

As for memory usage during alignment using 4 cores, minimap2 and uLTRA have similar memory footprints across the three simulated datasets, but uLTRA uses about 1.7-2.6 times more memory on the biological datasets (Table 3). deSALT uses the most memory on the simulated datasets and has a memory footprint similar to uLTRA on the ALZ dataset but uses lower memory on the SIRV and DROS. When parallelizing over 19 cores, the relative memory usage between minimap2 and uLTRA stays largely the same (Table S2). However, deSALT slightly decreases its relative memory footprint compared to minimap2 and uLTRA.

**Table 3.** Peak memory usage of alignment using 4 cores.

| Dataset | uLTRA | uLTRA_<br>mm2 | minimap2 | minimap2_GTF | deSALT | deSALT_G<br>TF |
| --- | --- | --- | --- | --- | --- | --- |
| ENS | 15Gb | 20Gb | 17Gb | 22Gb | 38Gb | 38Gb |
| SIM_ANN | 17Gb | 21Gb | 18Gb | 18Gb | 38Gb | 38Gb |
| SIM_NIC | 19Gb | 23Gb | 19Gb | 19Gb | 39Gb | 39Gb |
| SIRV | 6Gb | 8Gb | 3Gb | 3Gb | 2Gb | 2Gb |
| ALZ | 52Gb | 64Gb | 20Gb | 19Gb | 50Gb | 40Gb |
| DROS | 12Gb | 17Gb | 7Gb | 6Gb | 13Gb | 13Gb |

For indexing, uLTRA and minimap2 are relatively fast, while deSALT is slower (Table 4). uLTRA used the smallest amount of memory (Table 5), which is not surprising as it is processing a smaller region of the genome.

**Table 4.** Runtime of indexing.

| Dataset | uLTRA | minimap2 | deSALT |
| --- | --- | --- | --- |
| HG38 | 23m | 5m | 2h 49m |
| SIRV | 0m | 0m | 1m |
| DROS | 3m | 0m | 9m |

**Table 5.** Peak memory usage of indexing.

| Dataset | uLTRA | minimap2 | deSALT |
| --- | --- | --- | --- |
| HG38 | 7Gb | 19Gb | 74Gb |
| SIRV | 0Gb | 0Gb | 2Gb |
| Drosophila | 3Gb | 2Gb | 6Gb |

## Discussion

Splice alignment is an algorithmic problem central for the detection and prediction and quantification of isoforms. We have presented a novel splice alignment algorithm, and its implementation uLTRA. uLTRA aligns long transcriptomic reads to a genome using an annotation of coding regions. In addition, uLTRA can also run as a wrapper around minimap2. In this mode, it refines alignments around gene regions. uLTRA outputs alignments in SAM-format, and classifies the splice alignments according to the classification given in (Tardaguila et al. 2018) under an optional tag in the SAM-file. We evaluated uLTRA on simulated, synthetic, and biological data, and our analysis highlights some of the challenges with splice alignment and the current state-of-the-art approaches.

Using simulated data, we demonstrated uLTRA’s increased accuracy over other aligners. Particularly, uLTRA outperformed other state-of-the-art splice aligners when aligning reads to small exons. We also observed that uLTRA had high accuracy when aligning reads that could come from alternative haplotype sequences and novel isoform structures (Fig. S2), while other methods had a substantial decrease in accuracy on this dataset.

We used synthetic data to investigate the performance of alignment algorithms when aligning reads to known splice sites. Our experiments demonstrate that uLTRA aligns a much higher percentage of reads to known isoforms in the data. This holds true when running uLTRA as a wrapper around minimap2, indicating that uLTRA’s alignments are preferred based on edit distance of the alignments. Furthermore, uLTRA’s FSM alignments are distributed across the 59 isoforms with distinct splice-sites without indication of alignment bias towards specific isoforms as other aligners have (Fig. S3).

On biological data we demonstrated several examples where uLTRA aligns reads to the correct isoform structure while the other aligners do not. We showed several examples where isoforms containing small exons were misaligned (Fig. S9, S11). We also illustrated that employing junction specific alignment penalties may lead to concordant but erroneous alignment around junctions (Fig. S10). Finally, we observed cases where homopolymer differences in reads may lead to subtle alignment differences causing alignment to novel junctions (Fig. S12). In summary, the examples we provide on biological data demonstrate that using simple concordance analysis between aligners to measure accuracy can be misleading. Furthermore, the examples (Fig. S9-S13) came from genes that have been studied or linked to Alzheimer’s disease with many of them highly abundant. As several of these isoforms may not be detected with other alignment software, we demonstrated the utility of uLTRA and highlighted the significance of further development of splice alignment techniques.

We observed a large fraction of reads in the biological dataset that came from genomic regions in *Homo Sapiens* and *Drosophila* that are two well-annotated genomes. The large majority of these reads were aligned as NO_SPLICE (Fig. 3) and are likely to be intra-priming artifacts produced by long-read protocols (Tardaguila et al. 2018). In such cases, having an aligner that is not limited to aligning to only gene regions is preferred, as uLTRA will either not align them (as is the case for most of these reads; Fig. 3), or worse, overfit them to gene regions. We observed that this is resolved by using uLTRA as a wrapper around minimap2. Overall, our experiments on simulated, synthetic, and biological data indicated that uLTRA_mm2 (i.e., uLTRA as a wrapper around minimap2) produced the most favorable alignments at the cost of an increased runtime.

As alignment to pan-genome graphs has demonstrated its advantage over linear genomes (Garrison et al. 2018), it is of interest to explore such approaches in a transcriptomic setting. Our alignment strategy facilitates the addition of variant sequences in a relatively straightforward manner by adding alternative segments to uLTRA’s index (containing variations obtained from variant annotation file). We, therefore, aim to continue working on uLTRA in this direction.

## Conclusion

We present a new splice alignment algorithm and its implementation uLTRA. Our method models splice alignment as a two-pass collinear chaining problem with a novel exon chaining formulation. Our analysis highlights some of the challenges with splice alignment and the current state-of-the-art approaches. We show that uLTRA substantially improves splice-alignment accuracy of long RNA-seq reads using biological, spike-in, and simulated datasets. Furthermore, uLTRA can be used both as a stand-alone aligner and as a wrapper around minimap2 to handle reads aligning to unannotated regions. Furthermore, we demonstrate several examples on biological data where uLTRA aligns reads to previously annotated and novel isoform structures that other aligners did not detect. This highlights the immediate utility that uLTRA has when profiling a new transcriptome.

## Methods

### Step 1: Indexing and processing the genome annotation

A *part* is defined as the smallest genomic region fully covering a set of overlapping exons (Fig. 1). By construction, parts are disjoint regions of the genome. *Flanks* are constructed by taking regions of size *F* nucleotides downstream and upstream of parts. If two parts are separated with a distance of less than *F* nucleotides, then the non-overlapping region between the two parts is chosen as a flank region (Fig. 1). By construction, flanks are disjoint regions, both to each other and to parts. Finally, segments are constructed from start and end coordinates of exons. Segments are constructed for each part individually as follows. For a sorted array of exon start and stop coordinates within a part, a segment is constructed for each pair (*x_i_*, *x*_*i*+1_) of adjacent coordinates in the array if *x*_*i*+1_ – *x_i_* ≥ *X* where *X* is a parameter to uLTRA (set to 25). If *x*_*i*+1_ – *x_i_* < *X*, uLTRA iteratively attempts to add segments in each direction until success. That is, uLTRA attempts to add (*x_i–k_*, *x*_*i*+1_) and (*x_i_*, *x*_*i*+1+*k*_) for *k* = 1, 2, …, until first success in each direction. Finally, there may be parts where *y* – *x* < *X* (see exon e7 in Fig. 1). Small segments, exons, or parts have a lower probability of containing a MEM (and therefore be omitted from alignment). We address this complication as follows. uLTRA stores all exons and segments smaller than a threshold in a container that links gene ID to the small segments. This data structure will be queried, and all small segments will be included, whenever there are MEMs to segments linked to the same gene ID.

### Step 2: Collinear chaining with MEMs

A *Maximal Exact Match* (*MEM*) ([a..b],[c..d]) means that genome segment [a..b] matches read segment [c..d], and that such match cannot be extended to either direction. We use notation A[i].x to denote the endpoints of MEMs for x ∈ {a,b,c,d}. Let array A[1..n] contain the MEMs. A *chain* S is a collinear subset of A, meaning that S[i].a<S[i+1].a and S[i].c<S[i+1].c for 0<i<n (i.e. satisfying the weak precedence (Mäkinen and Sahlin 2020)). Coverage(S) is defined as the number of identities in an alignment induced by S, i.e., the length of the anchor-restricted LCS (longest common subsequence) of reference and the read, where anchor now means a MEM (Mäkinen and Sahlin 2020): If there are no overlaps between MEMs in chain S, Coverage(S) is the overall length of MEMs in S, but if there are, the score is adjusted by adding only the minimum length of the non-overlapping parts of the consecutive MEM intervals (Mäkinen and Sahlin 2020). Here we look for chains that have no overlaps in the genome, so for finding S that maximizes Coverage(S), we use Algorithm 1 in (Mäkinen and Sahlin 2020) that runs in O(n log n) time.

### Step 3: Collinear chaining of MAMs

We refer to an *approximate match*, as an alignment of a genome segment [a..b] to a read segment [c..d] with an accuracy higher than a threshold (parameter to uLTRA). Here, accuracy is defined as the number of matches divided by the length of the alignment. We find approximate matches of the genome segment by aligning it in semi-global mode to the read using edlib. The length of the alignment is defined by the genome segment’s first and last nucleotide coordinates. A *Maximal Approximate Match* (*MAM*) ([a..b],[c..d]) means that genome segment [a..b] matches approximately read segment [c..d] and that no other approximate match has higher accuracy on the read. Furthermore, we let λ ∈ [0,1] be the penalty for each nucleotide that overlap (on the read) between two MAMs and δ ∈ [0,1] the penalty for the distance between two MAMs (on the read). Let array *A*[1, …, *N*] contain the MAMs where we use the following notation: *A*[*i*]. *a,A*[*i*]. *b*, *A*[*i*]. *c,A*[*i*]. *d A*[*i*]. *acc* to denote the genome start, genome stop, read start, read stop, and accuracy of MAM *i*. Let *S* [1, …, *m*] be a chain of the MAMs in *A* under the weak precedence constraint (Mäkinen and Sahlin 2020). For two MAMs *x, y* in *A*, we introduce the following functions. Let *v*(*x*) = (*x. d* – *x. c*), *o*(*x,y*) = *max*{0, *x.d* – *y. c*} (the overlap), and *d*(*x,y*) = max {0, *y.c* – *x.d*}(the distance between MAMs) on the read, then the *score*(*S*) of a MAM-chain is defined as

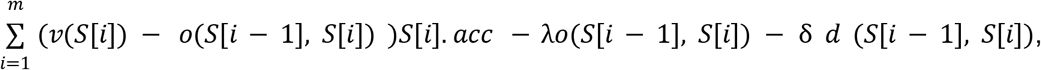

where *o*(*S*[0], *S*[1]) = *d*(*S*[0], *S*[1]) = 0

We find the chain 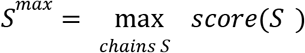. This formulation intuitively selects the solution with the best coverage and accuracy, while penalizing overlapping MAMs or MAMs that occur far apart. This formulation is solved with a dynamic programming algorithm: Sort array *A*[1, …, *N*]by values *A*[*i*]. *a*. Let *W*[0, …, *N*] be the target array, where we wish to store for each *W*[*i*] the maximum score over chains ending at MAM *A*[*i*]. To compute *W*[*i*], one can consider adding *A*[*i*] to chains ending at *A*[*i*’], ∀*i*’ < *i* with *A*[*i*’]. *c* < *A*[*i*]. *c*. This increases the score by *w*(*i*’, *i*) = (*v*(*A*[*i*]) – *o*(*A*[*i*’], *A*[*i*]))*A*[*i*]. *acc* – λ *o*(*A*[*i*’], *A*[*i*]) – δ *d* (*A*[*i*’], *A*[*i*]). After initializing *W*[0] = 0, we can set *W*[*i*] to the maximum over *W*[*i*’] + *w*(*i*’, *i*) for 0 ≤ *i*’ < *i* with *A*[*i*’]. *c* < *A*[*i*]. *c* from left to right, and the maximum scoring chain can be traced back starting from the maximum value in *W*[1, …, *N*]. Although this computation takes quadratic time, in practice the instances of segments are small enough to be solved quickly. It is not known whether our formulation allowing weighted hits, overlap, and gap penalties can be solved in subquadratic time, although recent breakthroughs have been made for chaining problems allowing overlap and gap costs (Jain, Gibney, and Thankachan, 2021).

### Wrapping around minimap2

uLTRA can be used as a wrapper around minimap2 to detect alignments outside annotated regions. In this mode, uLTRA first runs minimap2. After reads have been aligned with minimap2, uLTRA parses minimap2’s alignments to find reads with primary alignments outside the regions indexed by uLTRA. A read with more than a fraction of *X* nucleotides (parameter to uLTRA; *X* = 0.1 used here) out of the total aligned nucleotides is considered *genomic* and not realigned with uLTRA. We use an interval-tree data structure to hold the indexed regions to find overlap of a read and indexed regions. This permits an *O*(*log Q*) query time, where *Q* is the number of intervals on the chromosome to which the read is aligned. The alignments that are classified as genomic are not aligned with uLTRA. uLTRA then proceeds to align all remaining reads. Instead, uLTRA will report minimap2’s primary alignments for these reads. uLTRA then proceeds to align all reads not classified as genomic as described. In a final step, uLTRA compares the reads that have been aligned with both aligners and selects the best alignment based on edit distance to the genome. The final output SAM-file consists of the best alignments to uLTRA-indexed regions and minimap2’s alignments of genomic reads.

### Implementation

#### Chaining of MEMs

In the implementation, the optimal solution instance is found through backtracking. If several possible tracebacks paths lead to the same optimal value for a given optimal value in the traceback vector, uLTRA will always choose the closest MEM, i.e., the one with the highest index j. This means that the mem with the closest genomic coordinate is chosen if several exist.

If several optimal chaining solutions are found, i.e., several positions in the vector traceback vector have the optimal value, uLTRA will report all of the solutions by backtracking each instance (as described above). This is not a rare case since there can be identical or highly similar gene copies annotated on the genome that give the same optimal value.

Since each read can have several chaining instances to solve, uLTRA pre-calculates the theoretical maximum MEM coverage that an instance can have, which is upper bounded by the sum of all the regions covered by mems in the reads. uLTRA then solves the chaining instances by highest upper bound on coverage. If at any point the upper bound drops below a drop-off threshold (parameter to uLTRA) the current best solution uLTRA skips to calculate the rest of the instances. There is also a parameter to limit the number of reported alignments.

#### Chaining of MAMs

MAMs are formed by aligning segments and exons with at least an alignment identity of X% (default 60), and in case of exons between 5-8 nucleotides in length, an exact match is required. Exons of 4bp or less are ignored because of the potential blowup in the number of matches across the read. Similarly to the MEM chaining, the traceback will choose the MAM with the highest index j.

#### Alignment reporting

The exons that are included in an optimal solution of the MAM chaining are concatenated into an augmented reference, and the read is aligned to this reference using parasail in semi-global mode. The alignment score and cigar string are computed from the alignment. Among all MAM instances for a read, the highest scoring one is selected as the primary alignment. If a read has multiple best scoring alignments, the one with the shortest genomic span of the alignment is reported, and if still a tie, an FSM is preferred over the other read labels.

A read is assigned as unaligned if the alignment score is lower than X*m*r, where r is the read length, m is the match score (set to 2 in parasail; see Suppl. Note A for details), and X is a parameter to uLTRA (set to 0.5). The default setting roughly corresponds to classifying a read as unaligned if it has more than 25% errors, or if a larger segment of the read is from, e.g., from a region that is not included in the indexing.

#### Output

uLTRA outputs alignments in SAM-file format with genomic coordinates as annotated by the transcript database. In addition, uLTRA outputs an annotation of the alignment following the definitions in (Tardaguila et al. 2018) in the SAM-file in the optional field “CN”.

## Data access

The biological and synthetic datasets are publicly available datasets. The pacbio Alzheimer dataset can be downloaded at https://downloads.pacbcloud.com/public/dataset/Alzheimer2019_IsoSeq/. The drosophila ONT and SIRV datasets can be downloaded from ENA under project accession number PRJEB34849. The Ensembl cDNA was downloaded from https://www.ensembl.org/biomart/martview/. All scripts used for simulating datasets and to run the evaluation are found at https://github.com/ksahlin/ultra/tree/master/evaluation. The source code of uLTRA is available at https://github.com/ksahlin/ultra.

## Competing interests

The authors declare that they have no competing interests.

## Acknowledgements

We would like to thank Paul Medvedev for helpful feedback and comments. The computations were performed on resources provided by the Swedish National Infrastructure for Computing (SNIC) at Uppsala Multidisciplinary Center for Advanced Computational Science (UPPMAX).

## Authors’ contributions

Both authors participated in the initial conception of the project. K.S. designed the method with input from V.M. K.S. implemented the algorithm, designed the experiments and analyzed the results. K.S. drafted the paper with significant contributions by V.M. Both authors critically revised the text.

## Funding

The work was partially supported by the Academy of Finland (grant 309048).

## Notes

### Competing Interest Statement

The authors have declared no competing interest.

### Summary of Updates

Improved methods. uLTRA now uses less time and memory. uLTRA can also align to unannotated regions by wrapping around minimap2. This enables novel gene detection. Refined analysis.

