## Supplementary data for "Accurate spliced alignment of long RNA sequencing reads"

#### Note A: Alignment parameters

##### uLTRA

We ran uLTRA (v 0.0.2, commit 04290f6) for all five datasets.

###### Indexing

We ran indexing, which consists of two steps `prep_splicing` and `prep_seqs` as follows.

uLTRA prep_splicing --disable_infer GTF_annotation outfolder

uLTRA prep_seqs ref_fasta  outfolder --min_mem X

Where X=17 for DROS, SIRV, and the simulated read datasets, and X=20 for ALZ and ENS data. Also, the prep_splicing method sets the following default parameters: --flank_size 1000, --mask_threshold 200, --small_exon_threshold 200.

###### Alignment

Within the algorithm, uLTRA sets the --ont parameter for the DROS, SIM_ANN, and SIM_NIC datasets to encompass an error rate of 5-12%, and --isoseq parameter for the SIM_ENS and ALZ dataset for lower error rates. The --ont parameter sets minimum MEM size of 17 and MAM alignment minimum accuracy of 0.6, while  --isoseq sets minimum MEM size of 20 and MAM alignment minimum accuracy of 0.8. uLTRA uses slaMEM (Fernandes and Freitas 2014) to find MEMs, edlib (Šošic and Šikic 2017) to find MAMs, and parasail (Daily 2016) to perform final alignment to exons. We use alignment with parasail in semi-global mode with alignment penalties chosen as match:2, mismatch:-2, gap open: -3, gap extend: -1.

uLTRA also sets the following parameters related to the number of alignments to report --dropoff 0.95, --max_loc 5.

#### Minimap2

We ran minimap2 (H. Li 2018) (v2.17-r974-dirty, GitHub commit c9874e2). We used -k 14 for PacBio Alzheimer dataset and -k 13 for the other datasets to index the reference genomes. For alignment with minimap2, we specified parameters --eqx -t 19 -ax splice -k14 -G 500k for the PacBio Alzheimer dataset, and --eqx -t 19 -ax splice -k13 -w 5 -G 500k for the rest of the datasets. The -k and -w parameters were set to 13 and 5, respectively, to improve accuracy over default parameters. We observed a 0.25%, 0.5%, and 0.72% increase in the number of correct alignments over default parameters on SIM_ENS, SIM_ANN, and SIM_NIC. We also increased the -G parameter from the default value of 200,000 to 500,000 in all datasets except for SIRV. The parameter -G controls maximum intron length. We observed that minimap2 missed valid high-quality mapped isoforms in the ALZ dataset with introns larger than 200,000 (e.g., 10 reads that uLTRA found to transcript ENST00000555571.5 with an intron of length 387,219nt).

Finally, minimap2 sets a special penalty score for non-canonical junctions. We set the non-default parameters --splice-flank=no --secondary=no -C5 on the SIRV dataset as suggested in the minimap2 documentation and discussed in issue 99 in the minimap2 repository since SIRV isoforms do not honor the canonical AG-GT junction to the same extent as biological data. We observed a drastic improvement with the setting --splice-flank=no compared to default splicing mode. With original splicing parameters, minimap2 only mapped reads as FSM to 45 unique isoforms.

#### deSALT

We ran deSALT and deSALT_GTF (v1.5.6, commit 733831b) on all datasets besides the ALZ dataset with deSALT_GTF for which the v1.5.6 version returned an error. On this dataset, we used an earlier version (v1.5.5). We indexed the genomes with default parameters, as suggested in (Liu et al. 2019). We ran deSALT alignment with parameters -d 10 and -s 2 for all datasets, as (Liu et al. 2019) used these settings for both ONT and PacBio Iso-Seq data. We used the parameter -l 14, which lowers the seeding kmer size from the default of 15 and is supposed to increase accuracy at the cost of runtime. We also increased the --max-intron-len parameter from the default value of 200,000 to 500,000 in all datasets except for SIRV. The parameter --max-intron-len controls maximum intron length, and we observed, similarly to minimap2, that deSALT missed valid isoforms with intron larger than the default value of 200,000.

Similar to minimap2, deSALT sets an individual penalty score for non-canonical junctions. We set the parameter --noncan to 4 instead of the default of 9 on the SIRV data, and, similarly to minimap2's results, and we observed a substantially improved accuracy over default parameters. With original splicing parameters, deSALT only mapped reads as FSM to 47 unique isoforms.

#### Graphmap2

We ran GraphMap2, github commit 11815ed. We could not install Graphmap2 on one Linux system and we received segmentation faults on another system. These issues were reported (GitHub issues 17 and 18).

### Note B: Simulated data

#### Simulating from annotated transcripts

We considered using already existing read simulators (NanoSim (Yang et al. 2017), DeepSimulator (Y. Li et al. 2018), SimLord (Stöcker, Köster, and Rahmann 2016), and SNaReSim (Faucon et al.("SNaReSim: Synthetic Nanopore Read Simulator - IEEE Conference Publication" n.d.)) before implementing a transcriptomic read simulator. However, they are genomic read simulators and cannot easily be modified to simulate full-length transcript reads.

In our simulations, we simulate transcripts from 234,207 distinct ENSEMBL cDNA sequences from the human reference. We sample reads uniformly from the transcripts. Each sampled transcript is then subject to sequencing errors. At each position in the transcript, we enter an error state with a probability of 0.05. If the simulator enters an error state, it chooses a deletion state with a probability of 0.5, substitution with a probability of 0.3, and insertion with a probability 0.2. In the deletion and insertion states, the length of the insertion and deletion is simulated from a geometric distribution with a parameter of 0.5. This model gives an average error rate of 8.6%.

#### Simulating novel transcripts

We used the GTF annotations for this simulation. We select all genes with four or more exons that have non-overlapping genomic coordinates. For each transcript we simulate from such a gene, we include the first and last exon with probability 1, but any internal exon with a probability of 0.5. This probabilistic inclusion of exons gives isoforms. We check if the transcript matches an already annotated isoform. If not, we keep it in the simulation. To keep the number of transcripts equal to the annotated number of transcripts for each gene, we simulate as many novel transcripts as the gene had annotated transcripts (if the number of possible combinations permits).

##### Evaluation of simulated data

For the simulated data, we have the true genome annotations of each exon in the transcript. A read is therefore classified as *correct* if the read is aligned to all the correct exons, and all of the exon alignments have an offset of fewer than 15 nucleotides to true annotation coordinates. An alignment is *inexact* if the read is aligned to the correct exons, but at least one junction offset is more than 15 nucleotides. An alignment is classified as having an *exon difference* if the read alignment is missing one or more exons, or a segment of at least 15 nucleotides is aligned to a genomic location not included in the set of true exons (i.e., appearing as a false exon). An alignment is classified as an *incorrect location* if it aligns to a genomic location not overlapping with the correct annotation. In our classifications above, we chose 15bp as the threshold because we did not observe deletions or insertions longer than 15 nucleotides in the simulated data. Therefore, an offset larger than 15 nucleotides indicates a misalignment rather than an actual deletion or insertion of this size in the read, causing a junction offset.

### Note C: SIRV analysis

The SIRV dataset was downloaded from ENA under project accession number PRJEB34849. This dataset was processed with pychopper (https://github.com/nanoporetech/pychopper, commit 6dca13d) which detects reads covering the entire transcript (full-length reads) by identifying barcodes present in the reads. Pychopper was run with default parameters.

The SIRV dataset consists of 68 synthetic transcripts from 7 different loci sequenced with ONT R9 technology (see (Sahlin and Medvedev 2021) for details). The transcripts from each locus differ in their splicing pattern. Eight out of the 68 transcripts contain only one exon, and therefore do not have a splice site. Furthermore, two isoforms SIRV701 and SIRV705 have identical splice sites and differ only in a 2nt offset in both the transcription start and stop site. Therefore we used 59 isoforms with distinct splice sites to investigate alignment performance around splice sites. The ONT SIRV dataset was constructed so that the isoforms should have roughly the same abundance in the sample (the SIRV E0 mix). While sequencing depth bias can cause significant differences in read depth due to the isoforms having different lengths, with the given depth of 1.4 million reads, we expect all the isoforms to occur in the sample.

### Note D: Biological datasets processing

The ALZ dataset was downloaded from https://downloads.pacbcloud.com/public/dataset/Alzheimer2019_IsoSeq/. This dataset has been processed with the standard Iso-Seq bioinformatics pipeline (SMRTlink 8.0 "IsoSeq" protocol) to contain only full-length reads. The DROS dataset was downloaded from ENA under project accession number PRJEB34849. This dataset was processed with pychopper (https://github.com/nanoporetech/pychopper, commit 6dca13d) using default parameters to identify full-length reads.

### Supplementary Figures

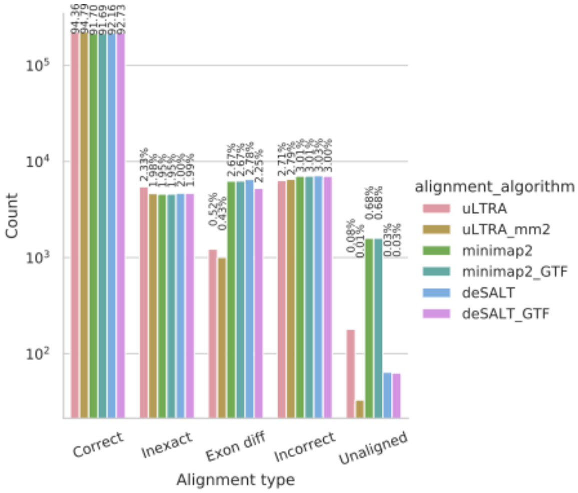


A

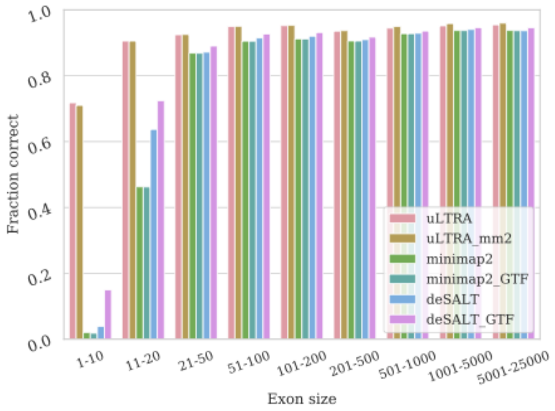


B

**Figure S1.** Alignment results on simulated data for the ENS dataset. (**A**) Percentage of reads in each respective category. (**B**) The fraction of correctly aligned exons (y-axis) as a function of exon size (x-axis).

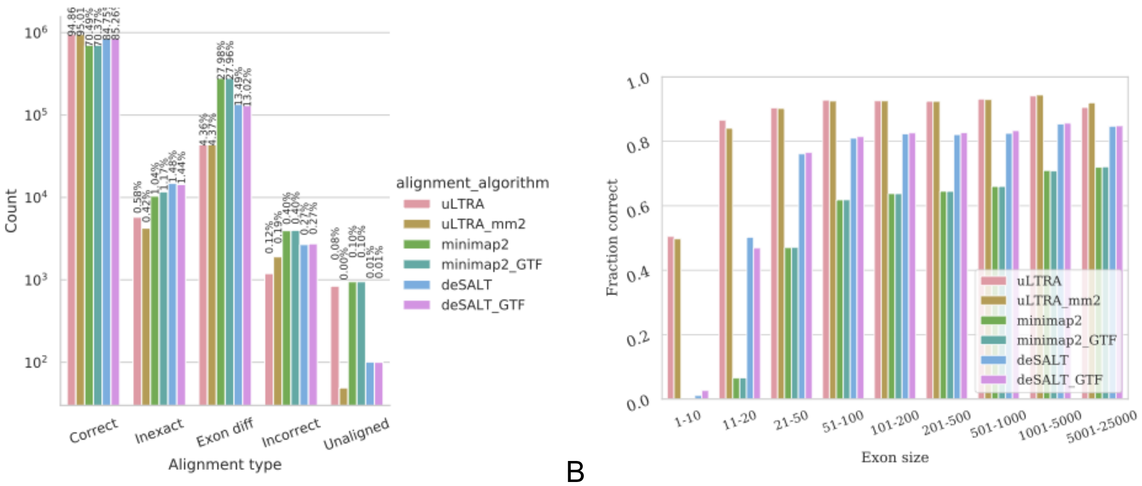


A B

**Figure S2.** Alignment results on simulated data for the SIM_NIC dataset. (**A**) Percentage of reads in each respective category. (**B**) The fraction of correctly aligned exons (y-axis) as a function of exon size (x-axis).

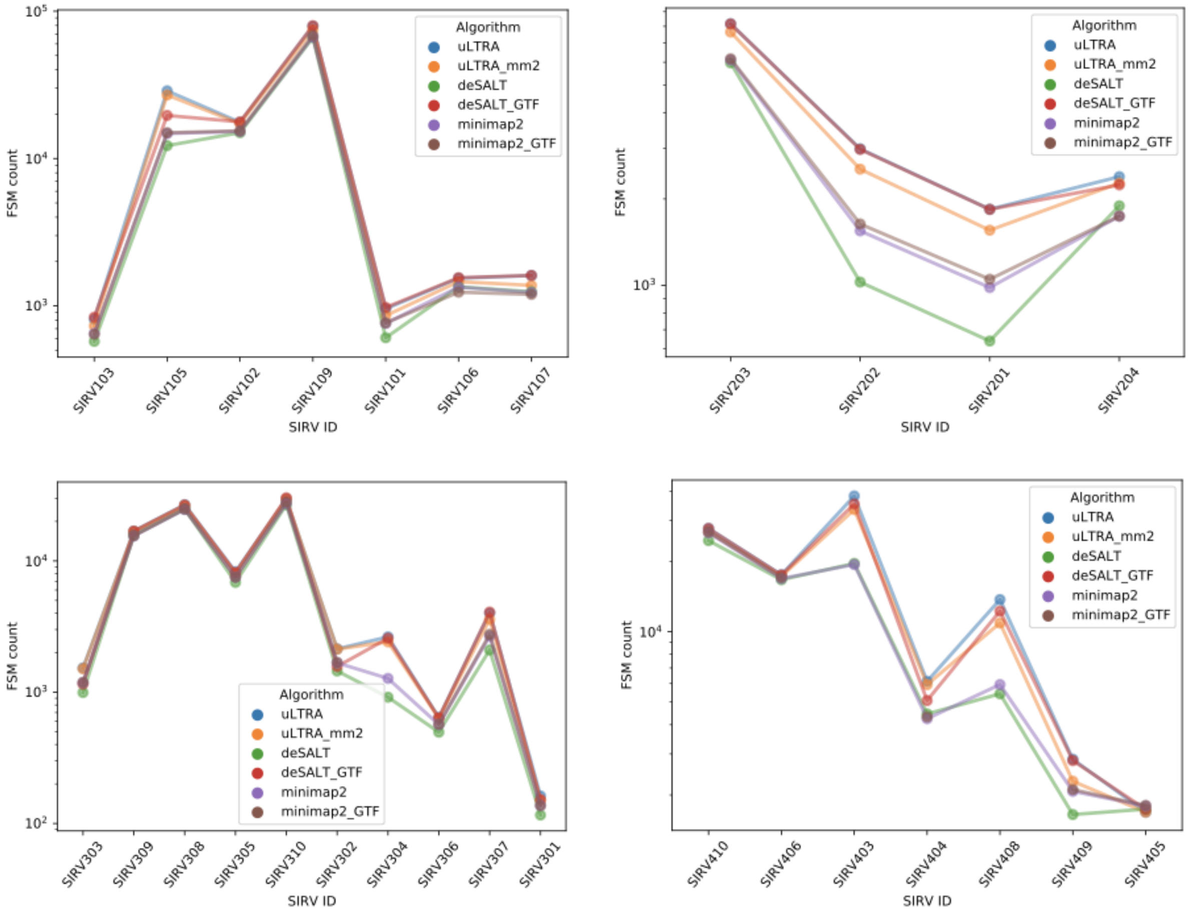

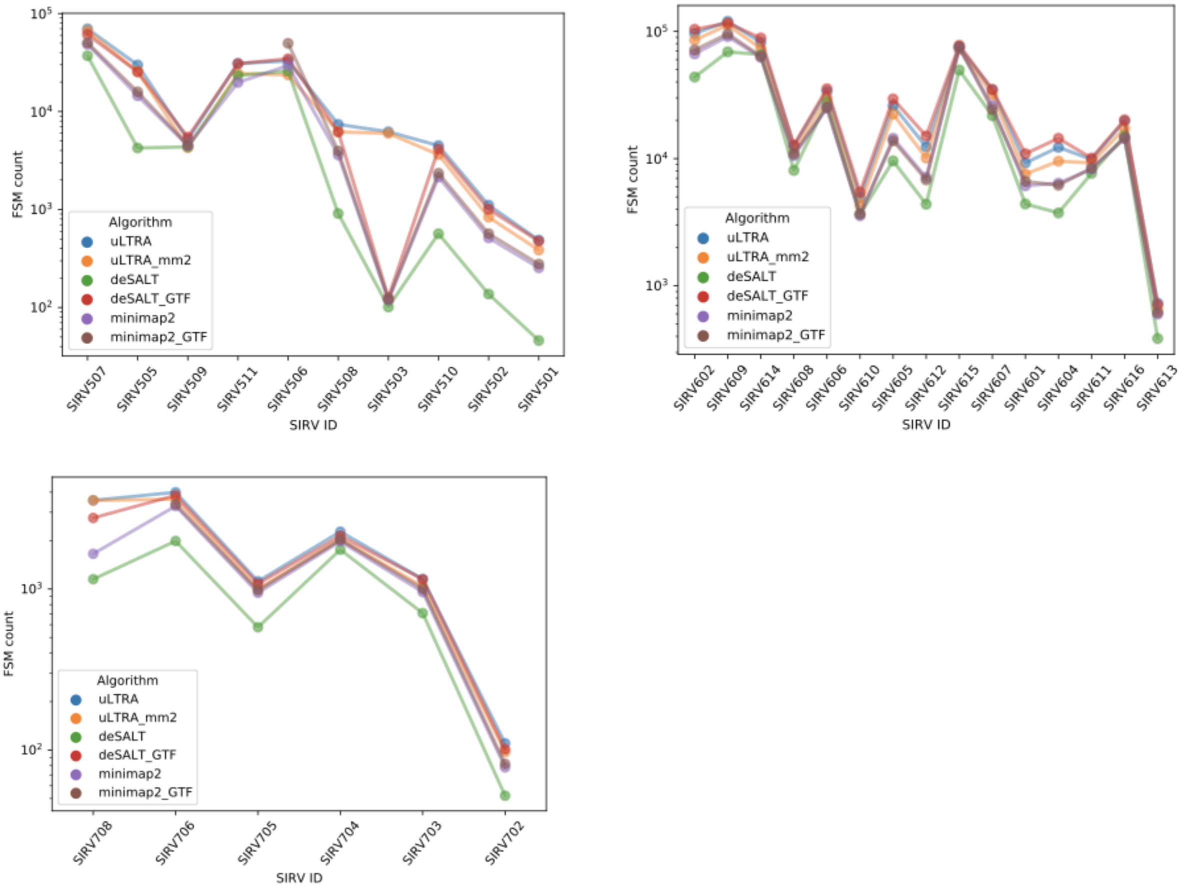


**Figure S3.** The number of reads annotated as FSM to each of the 59 SIRV isoforms with at least one splice site (y-axis log scale). One panel per gene loci.

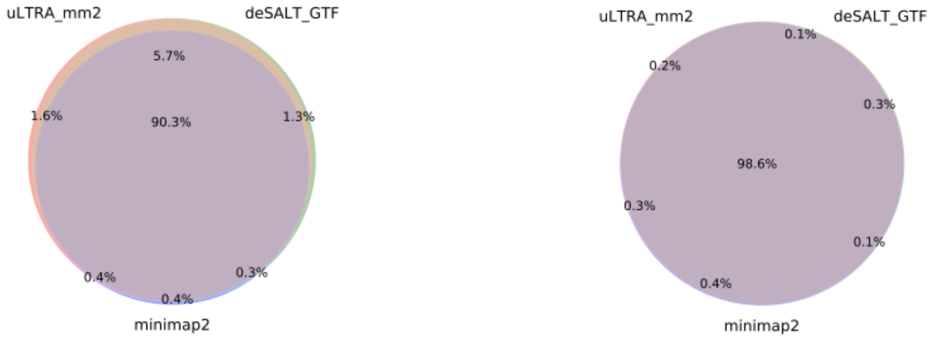


**Figure S4.** Alignment overlap concordance between uLTRA_mm2, deSALT_GTF, and minimap2 for DROS (A) and ALZ (B).

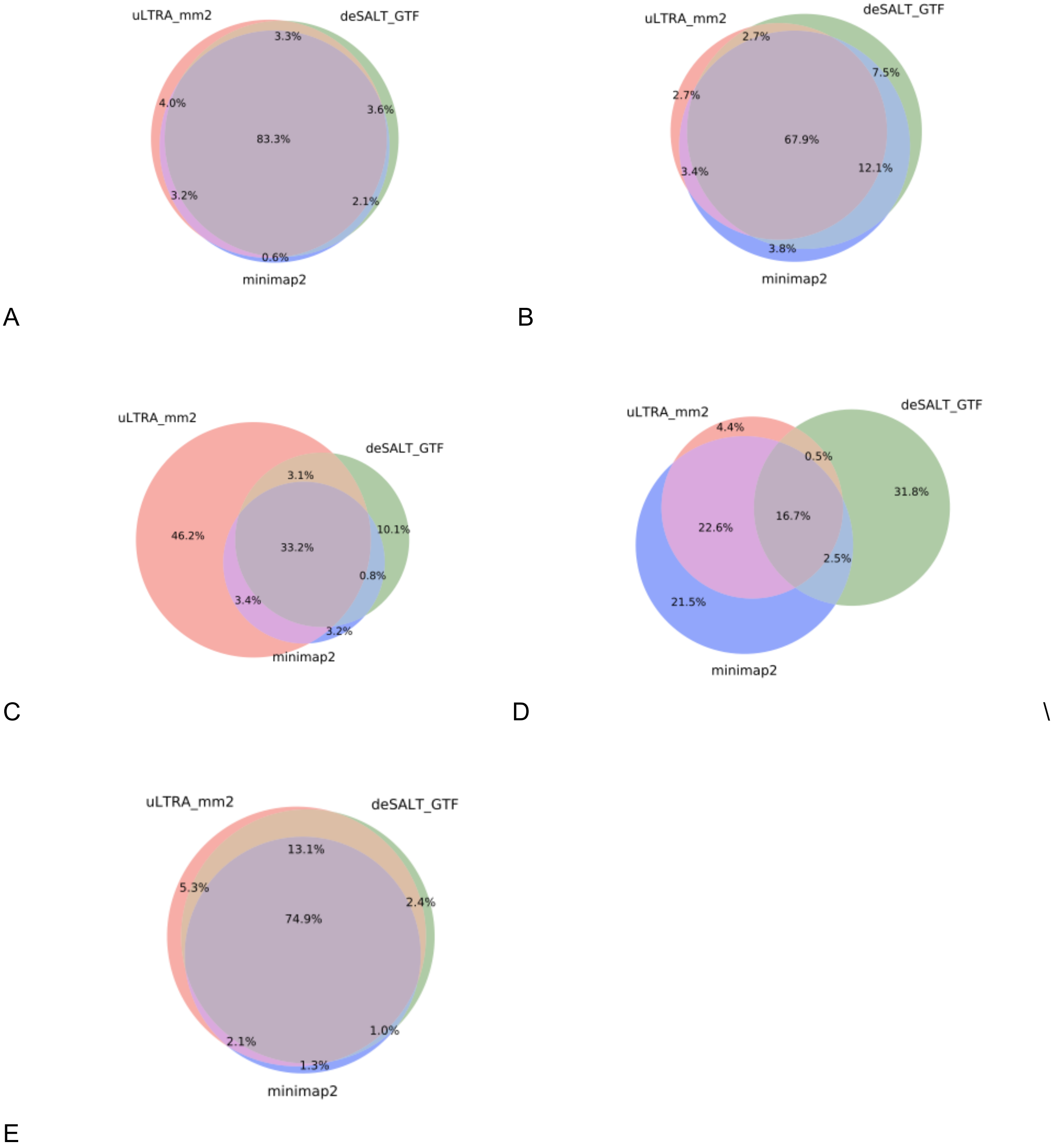


**Figure S5.** Alignment concordance between uLTRA_mm2, deSALT_GTF, and minimap2 for the different categories FSM (A), ISM (B), NIC (C), NNC (D) and NO_SPLICE (E) in DROS.

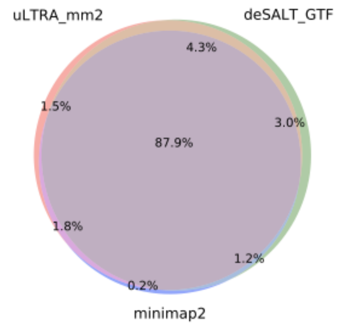


A

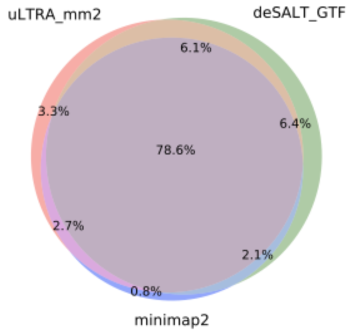


B

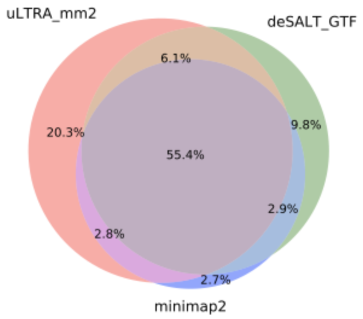


C

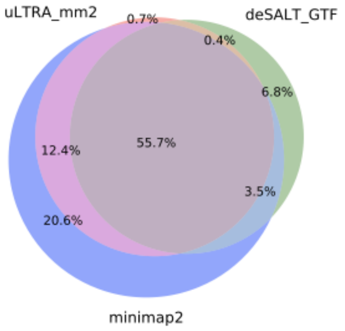


D

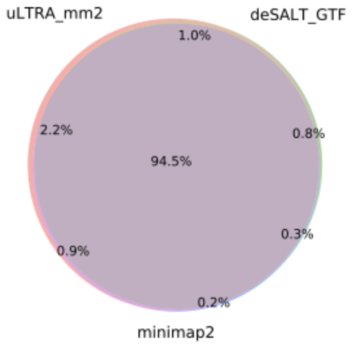


E

**Figure S6.** Alignment concordance between uLTRA_mm2, deSALT_GTF, and minimap2 for the different categories FSM (A), ISM (B), NIC (C), NNC (D) and NO_SPLICE (E) in ALZ.

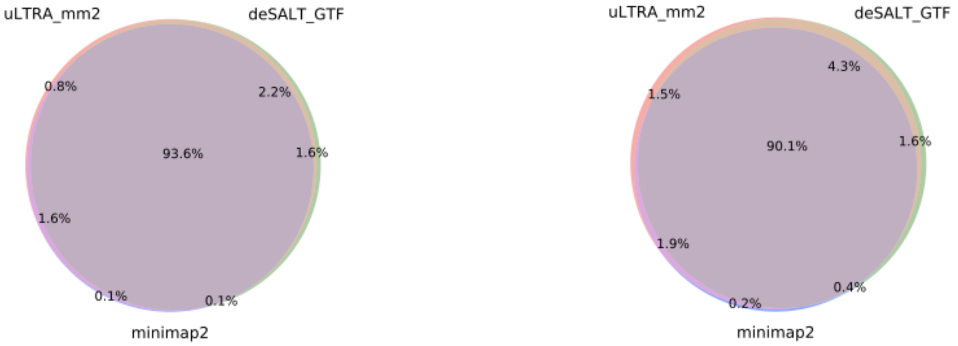


**Figure S7.** Concordance of unique isoforms that had FSM aligned reads for DROS (A) and ALZ (B).

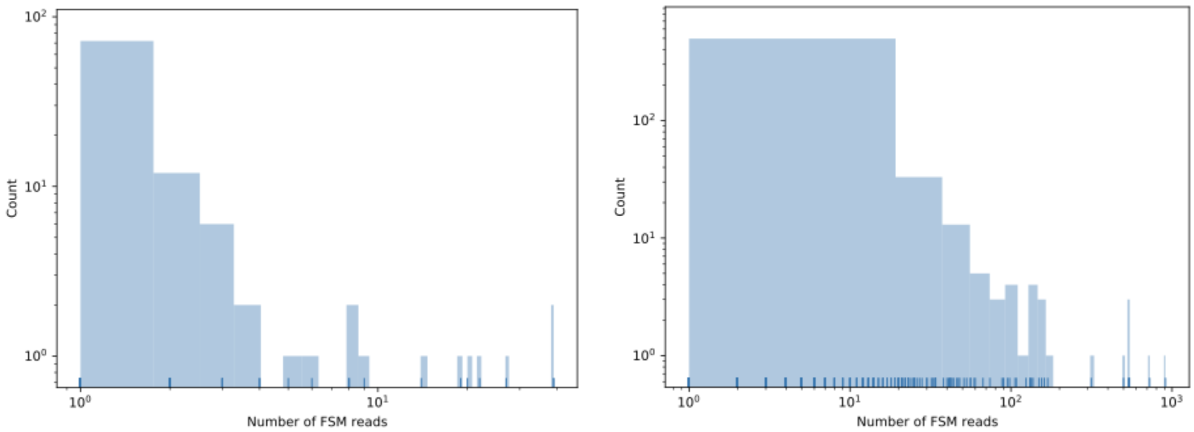


**Figure S8.** Histogram of the number of distinct isoform structures (y-axis) for which uLTRA_mm2 aligned FSM reads to, but deSALT_GTF and minimap2 did not. The x-axis shows the FSM read support for these isoforms. Panel (A) shows the DROS dataset and (B) the ALZ dataset.

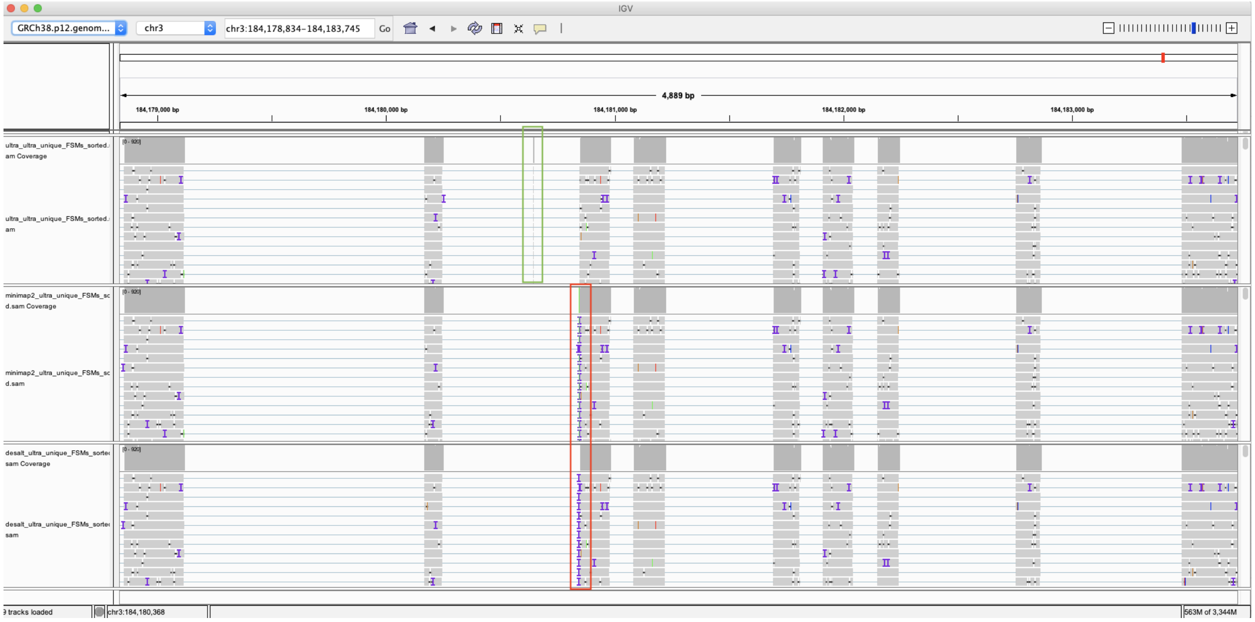
IGV
GRCh38.p12.genom...
chr3
chr3:184,178,834-184,183,745
Go
4,889 bp
184,179,000 bp
184,180,000 bp
184,181,000 bp
184,182,000 bp
184,183,000 bp
ultra_ultra_unique_FSMs_sorted.sam Coverage
ultra_ultra_unique_FSMs_sorted.sam
minimap2_ultra_unique_FSMs_sorted.sam Coverage
minimap2_ultra_unique_FSMs_sorted.sam
desalt_ultra_unique_FSMs_sorted.sam Coverage
desalt_ultra_unique_FSMs_sorted.sam
9 tracks loaded
chr3:184,180,368
563M of 3,344M

A

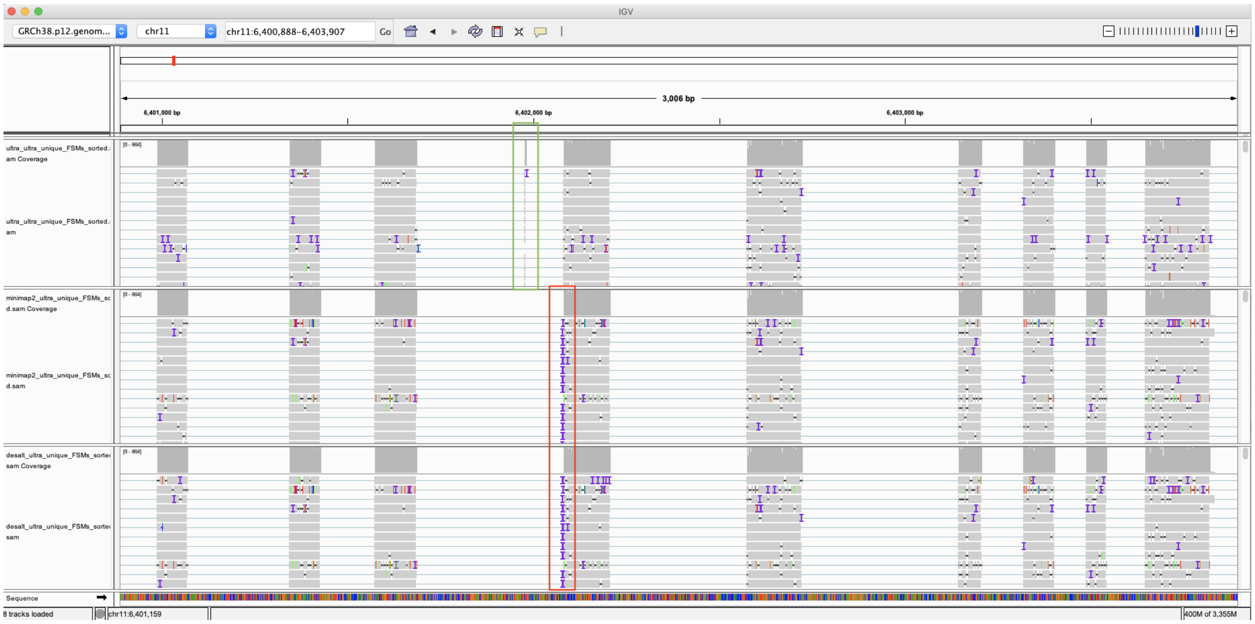
IGV
GRCh38.p12.genom...
chr11
chr11:6,400,888-6,403,907
Go
3,006 bp
6,401,000 bp
6,402,000 bp
6,403,000 bp
ultra_ultra_unique_FSMs_sorted.sam Coverage
ultra_ultra_unique_FSMs_sorted.sam
minimap2_ultra_unique_FSMs_sorted.sam Coverage
minimap2_ultra_unique_FSMs_sorted.sam
desalt_ultra_unique_FSMs_sorted.sam Coverage
desalt_ultra_unique_FSMs_sorted.sam
Sequence
8 tracks loaded
chr11:6,401,159
400M of 3,355M

B

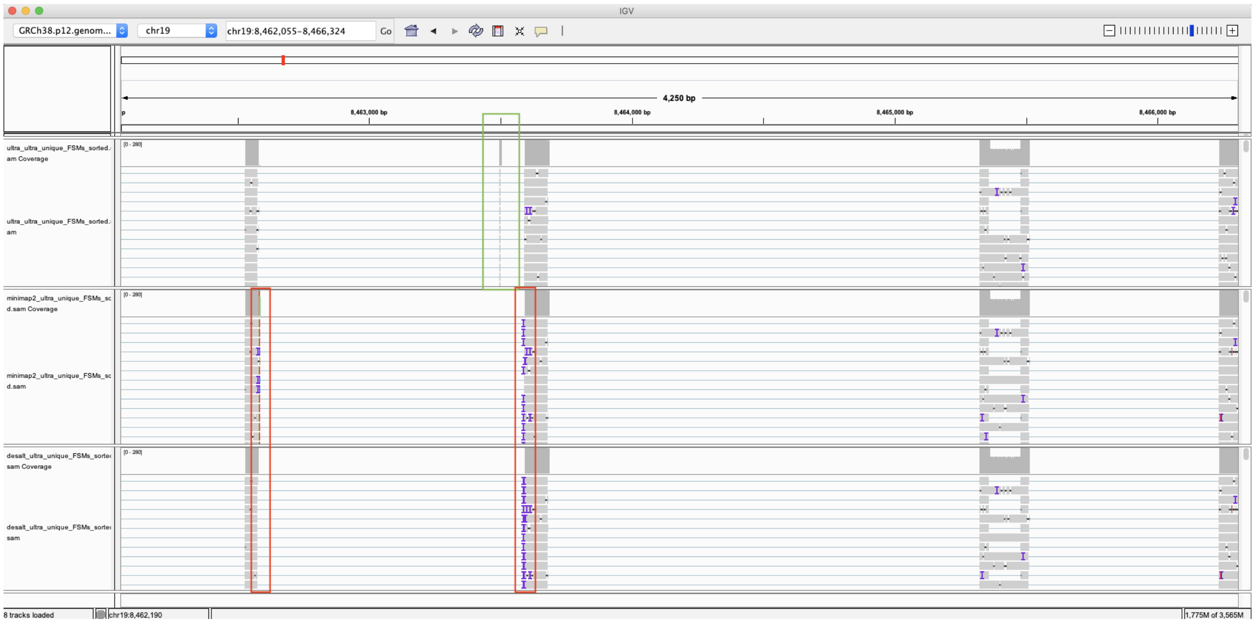


C

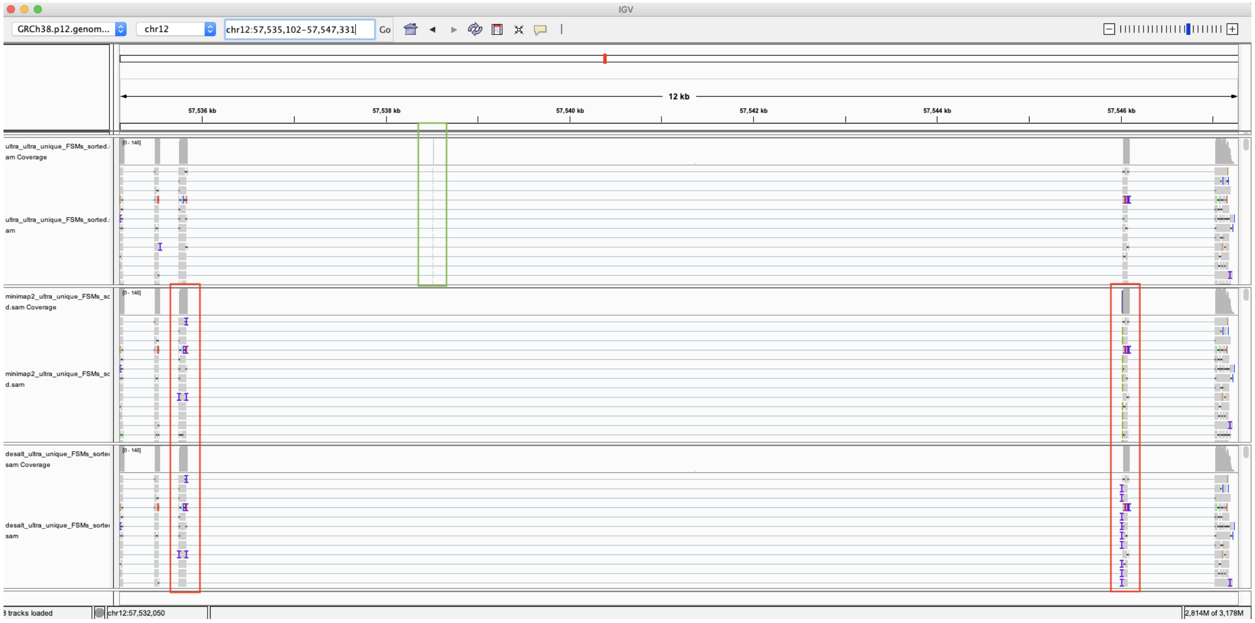


D

**Figure S9.** IGV tracks of FSM splice read alignments to unique FSM isoforms found by uLTRA_mm2 (first track) by aligning to small exons (6-8nt), with corresponding aligned reads of minimap2 (second track) and deSALT_GTF (third track). The green box highlights the best fit exon alignment. The red box highlights misalignments around the junctions caused by the unaligned small exons. (A) 910 reads mapping to transcript ENST00000292807.9 (AP2 gene). (B) 726 reads mapping to transcript ENST00000609360.6 (APBB gene). (C) 155 reads mapping to ENST00000325495.9 (HNRNPM gene). (D) 134 reads mapping to ENST00000543672.5 (DCTN gene).

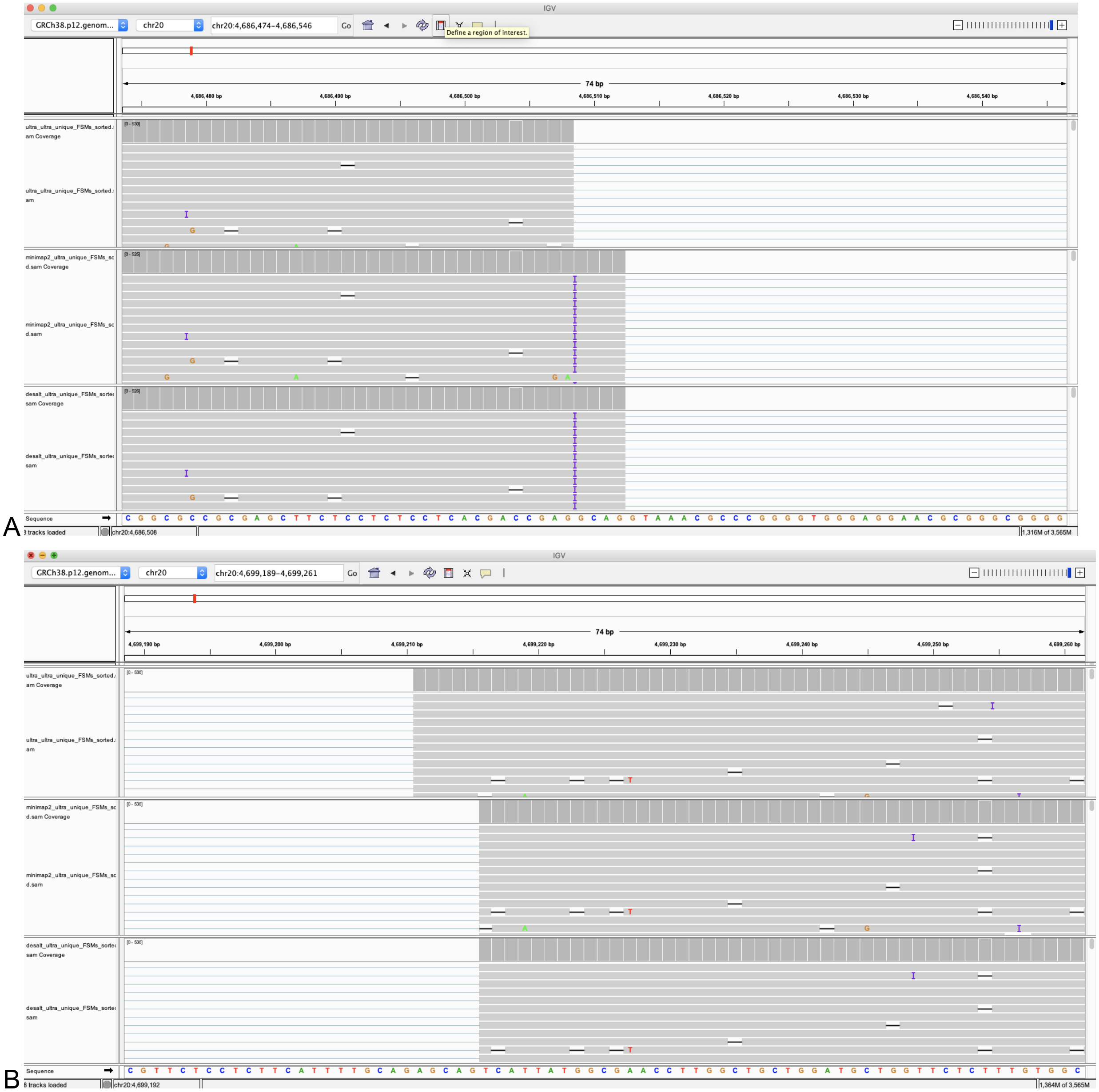


**Figure S10.** Potential subtle misalignment of 500 reads around a junction for minimap2 (second track) and deSALT_GTF (third track) cause structural change in splice junction. Both minimap2 and deSALT_GTF place a 5nt segment in the upstream junction (with a 1nt insertion) instead of placing the 5nt segment (full match) in the downstream junction (B). This is caused by overfitting the alignment to match a GT-AG junction with junction-specific penalties. In this gene, it appears as a small variant between the sample and the reference. Note that since the correct annotation is unknown for biological data, hence this may be a correct variation. However, uLTRA_mm2 achieves a higher identity in its alignment over the junction.

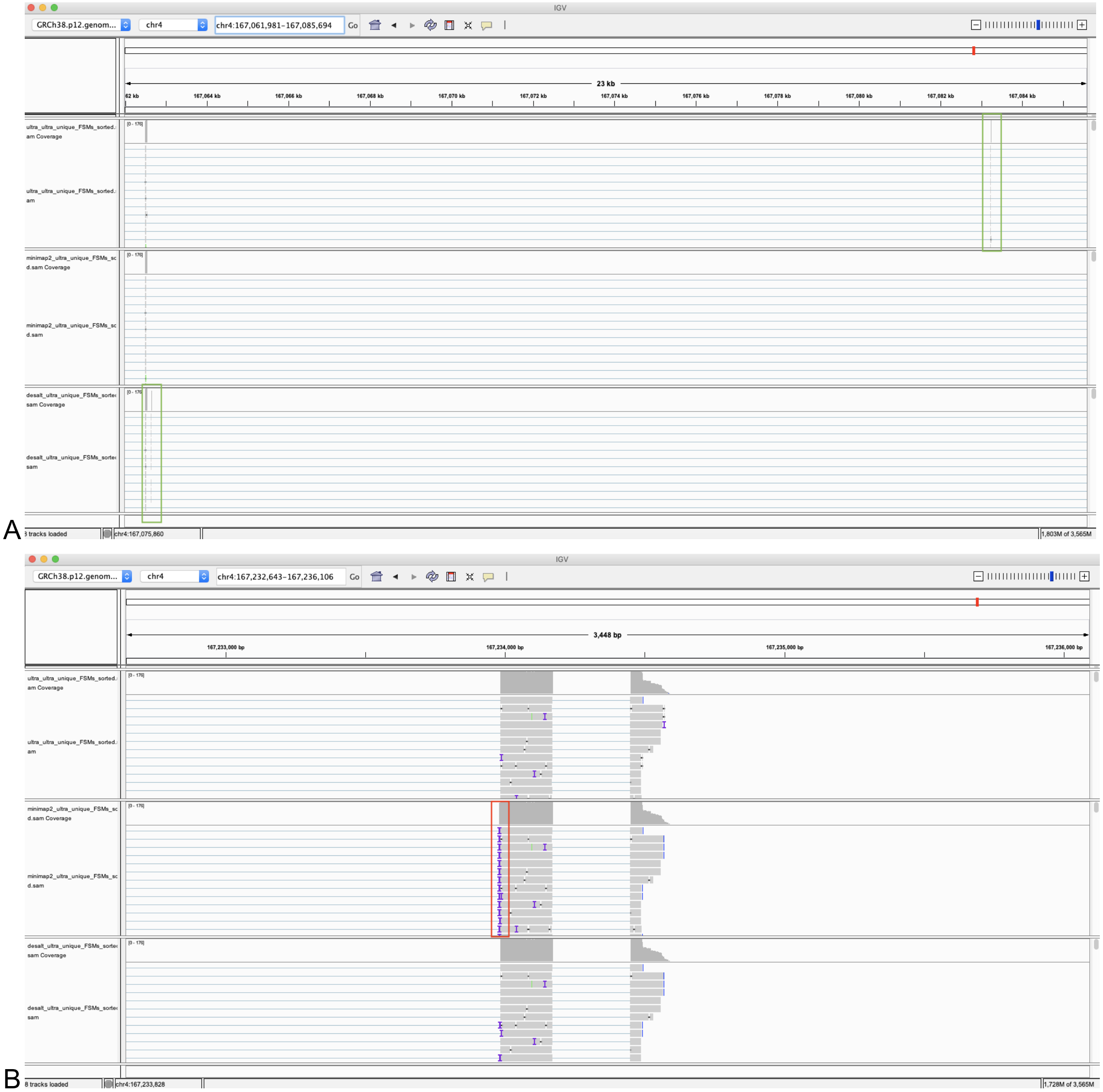


**Figure S11.** Discordant alignments of 161 reads between uLTRA (top track), minimap2 (middle track) and deSALT_GTF (bottom track) to the SPOCK gene. For these reads, uLTRA_mm2 and deSALT_GTF align a 9nt portion of the reads to two different exons (A) while minimap2 does not align this region and is instead present as an insertion in the downstream exon (B). uLTRA_mm2 chooses the upstream exon because of the deterministic implementation of taking the closest segment to the downstream hit in the traceback vector of the collinear MAM-chaining solution.

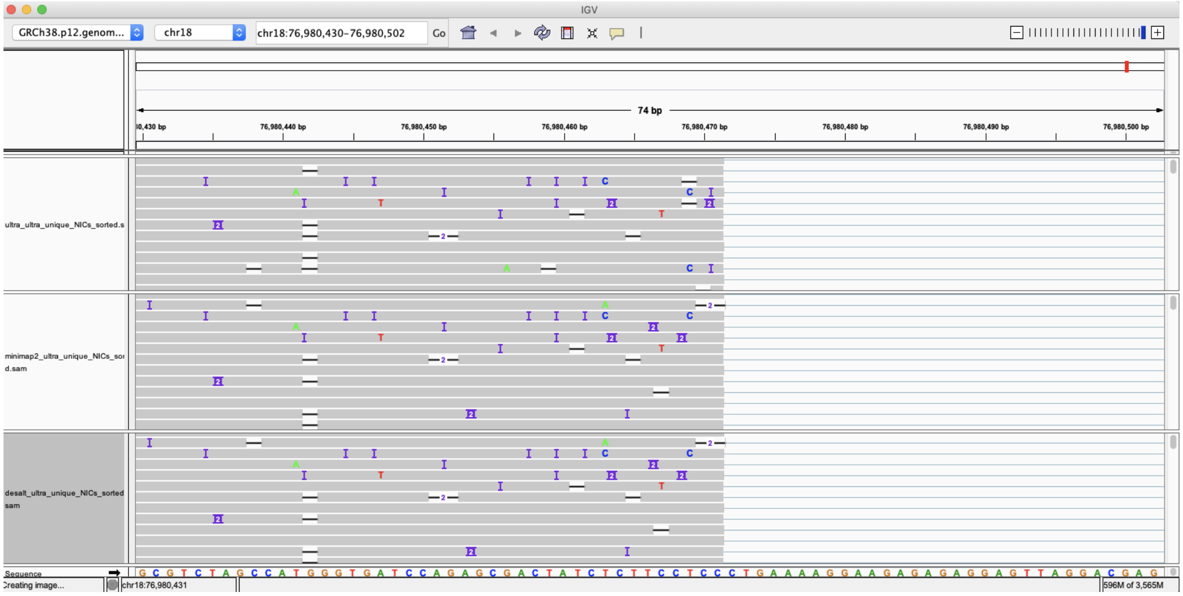


A

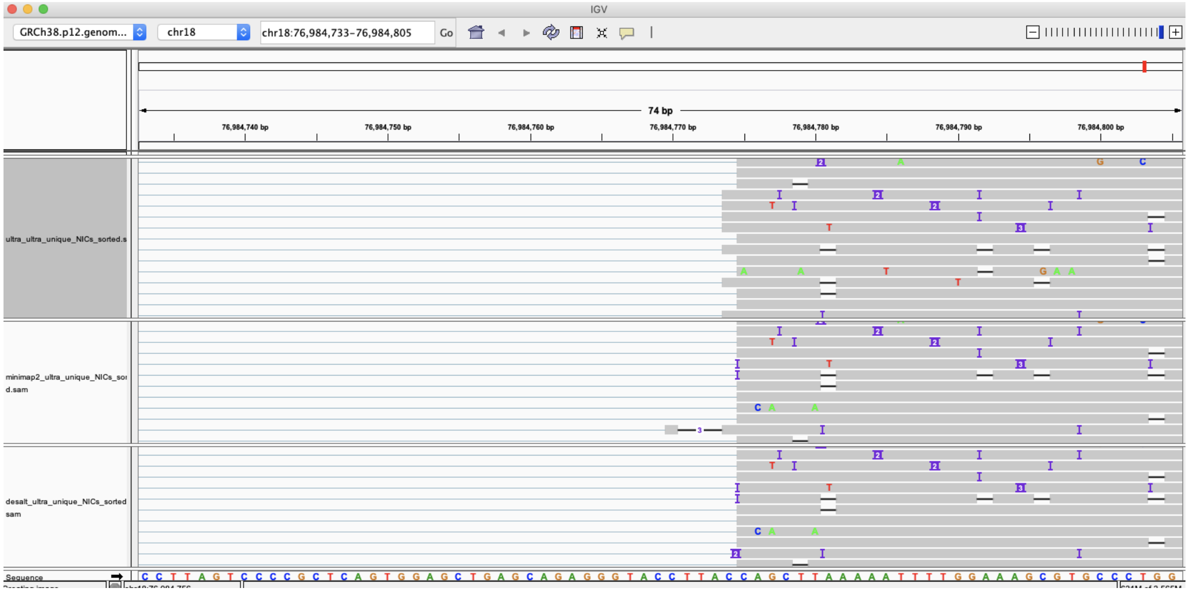


B

**Figure S12.** IGV tracks of splice read alignments to a junction of the MBP gene. Homopolymer stretches of C both upstream (A) and downstream (B) of the junction creates ambiguity in alignment. deSALT_GTF and minimap2 always align to the CT-AC junction by creating insertions of C at downstream junctions if needed, while uLTRA chooses a CT-TA junction for the reads with a homopolymer stretch of four Cs. The CT-AC matches an FSM transcript while the CT-TA junction creates a NIC transcript (943 reads).

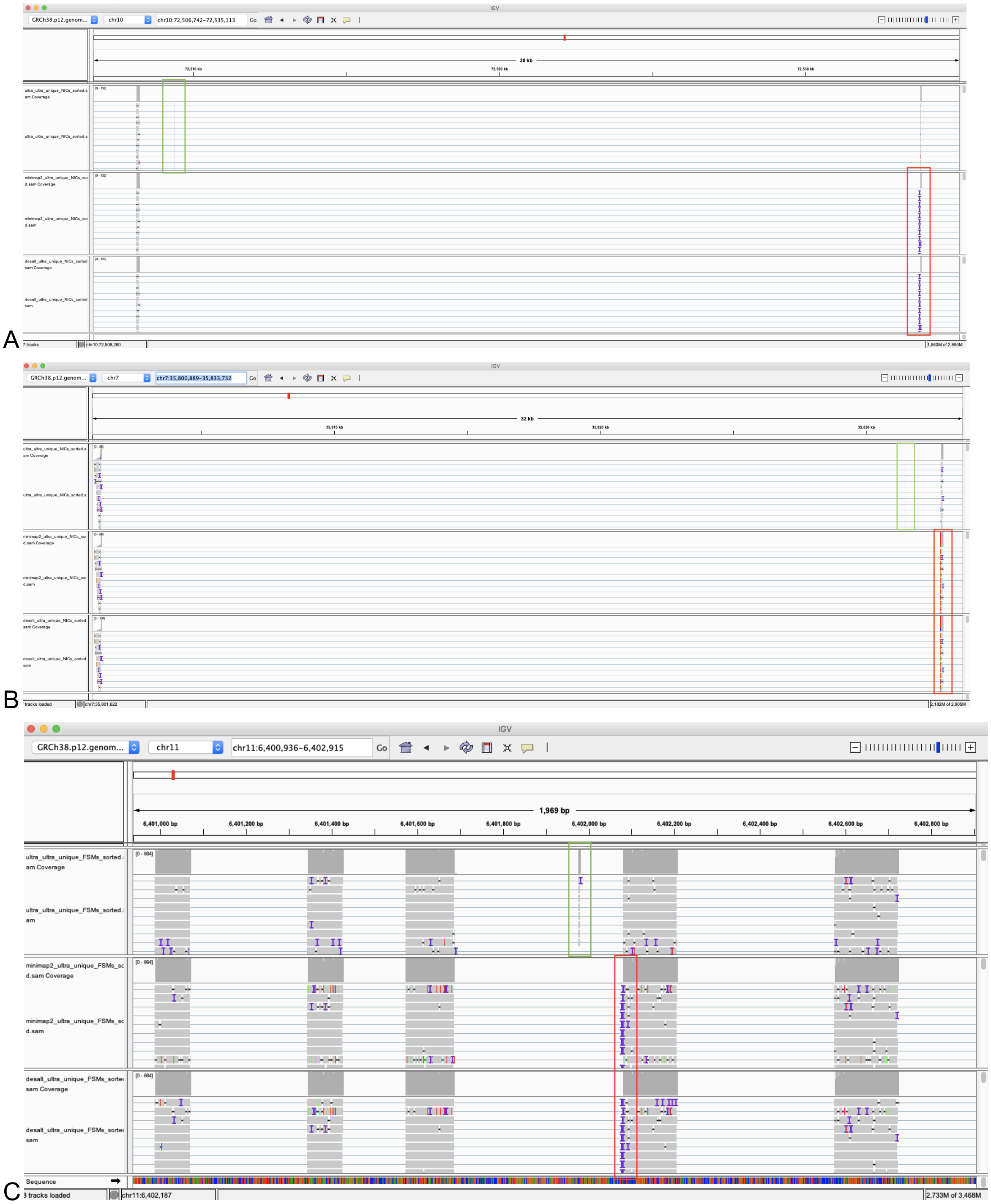
A
B
C
IGV
GRCh38.p12.genom...
chr11
chr11:6,400,936-6,402,915
Go
1,969 bp
6,401,000 bp
6,401,200 bp
6,401,400 bp
6,401,600 bp
6,401,800 bp
6,402,000 bp
6,402,200 bp
6,402,400 bp
6,402,600 bp
6,402,800 bp
ultra_ultra_unique_FSMs_sorted.
am Coverage
ultra_ultra_unique_FSMs_sorted.
am
minimap2_ultra_unique_FSMs_sc
d.sam Coverage
minimap2_ultra_unique_FSMs_sc
d.sam
desalt_ultra_unique_FSMs_sortec
sam Coverage
desalt_ultra_unique_FSMs_sortec
sam
[0 - 864]
Sequence
3 tracks loaded
chr11:6,402,187
2,733M of 3,468M

**Figure S13.** IGV tracks of splice read alignments to unique NIC isoforms found by uLTRA_mm2 (first track) by aligning to small exons (5-6nt), with corresponding aligned reads of minimap2 (second track) and deSALT_GTF (third track). A green rectangle highlights the best fit exon alignments. The red box highlights misalignments around the junctions caused by the unaligned small exons. (A) 126 reads mapping to a NIC transcript (with a 6nt exon) from the MICU1 gene. (B) 81 reads mapping to a NIC (with a 5nt exon) transcript of the SEPTIN7 gene. (C) 25 reads mapping to a NIC transcript (with a 6nt exon) from the APBB1 gene.

### Supplementary Tables

| Dataset | uLTRA | uLTRA_mm2 | minimap2 | minimap2_GTF | deSALT | deSALT_GTF |
|---|---|---|---|---|---|---|
| ENS | 51m | 51m | 13m | 14m | 6m | 6m |
| SIM_ANN | 1h 21m | 1h 45m | 45m | 47m | 23m | 23m |
| SIM_NIC | 1h 31m | 2h 41m | 1h 17m | 1h 19m | 30m | 32m |
| SIRV | 17m | 19m | 4m | 4m | 2m | 2m |
| ALZ | 4h 49m | 5h 42m | 2h 36m | 2h 48m | 2h 42m | 2h 41m |
| DROS | 43m | 42m | 6m | 7m | 8m | 8m |

**Table S1.** Runtime of alignment using 19 cores.

| Dataset | uLTRA | uLTRA_mm2 | minimap2 | minimap2_GTF | deSALT | deSALT_GTF |
|---|---|---|---|---|---|---|
| ENS | 69Gb | 70Gb | 28Gb | 28Gb | 38Gb | 38Gb |
| SIM_ANN | 70Gb | 74Gb | 29Gb | 30Gb | 40Gb | 40Gb |
| SIM_NIC | 70Gb | 73Gb | 32Gb | 32Gb | 39Gb | 39Gb |
| SIRV | 6Gb | 8Gb | 5Gb | 5Gb | 3Gb | 4Gb |
| ALZ | 90Gb | 102Gb | 28Gb | 30Gb | 136Gb | 85Gb |
| DROS | 18Gb | 23Gb | 11Gb | 12Gb | 32Gb | 32Gb |

**Table S2.** Peak memory usage of alignment.

### References

Daily, Jeff. 2016. “Parasail: SIMD C Library for Global, Semi-Global, and Local Pairwise Sequence Alignments.” *BMC Bioinformatics* 17 (February): 81.

Fernandes, Francisco, and Ana T. Freitas. 2014. “slaMEM: Efficient Retrieval of Maximal Exact Matches Using a Sampled LCP Array.” *Bioinformatics* 30 (4): 464–71.

Li, Heng. 2018. “Minimap2: Pairwise Alignment for Nucleotide Sequences.” *Bioinformatics* 34 (18): 3094–3100.

Liu, Bo, Yadong Liu, Junyi Li, Hongzhe Guo, Tianyi Zang, and Yadong Wang. 2019. “deSALT: Fast and Accurate Long Transcriptomic Read Alignment with de Bruijn Graph-Based Index.” *Genome Biology* 20 (1): 274.

Li, Yu, Renmin Han, Chongwei Bi, Mo Li, Sheng Wang, and Xin Gao. 2018. “DeepSimulator: A Deep Simulator for Nanopore Sequencing.” *Bioinformatics* 34 (17): 2899–2908.

Sahlin, Kristoffer, and Paul Medvedev. 2021. “Error Correction Enables Use of Oxford Nanopore Technology for Reference-Free Transcriptome Analysis.” *Nature Communications*. https://doi.org/10.1038/s41467-020-20340-8.

“SNaReSim: Synthetic Nanopore Read Simulator - IEEE Conference Publication.” n.d. Accessed June 21, 2019. https://ieeexplore.ieee.org/document/8031171.

Šošic, Martin, and Mile Šikic. 2017. “Edlib: A C/C ++ Library for Fast, Exact Sequence Alignment Using Edit Distance.” *Bioinformatics* 33 (9): 1394–95.

Stöcker, Bianca K., Johannes Köster, and Sven Rahmann. 2016. “SimLoRD: Simulation of Long Read Data.” *Bioinformatics*. https://doi.org/10.1093/bioinformatics/btw286.

Yang, Chen, Justin Chu, René L. Warren, and Inanç Birol. 2017. “NanoSim: Nanopore Sequence Read Simulator Based on Statistical Characterization.” *GigaScience* 6 (4): 1–6.

Faucon, P. C., Balachandran, P., and Crook, S. "SNaReSim: Synthetic Nanopore Read Simulator," *2017 IEEE International Conference on Healthcare Informatics (ICHI)*, Park City, UT, 2017, pp. 338-344, doi: 10.1109/ICHI.2017.98.
